# Remote memory engrams are controlled by encoding-specific tau phosphorylation

**DOI:** 10.1101/2023.12.01.569663

**Authors:** Kristie Stefanoska, Emmanuel Prikas, Yijun Lin, Renée Kosonen, Arne Ittner

**Affiliations:** Laboratory for Molecular Dementia and Memory Research, Flinders Health and Medical Research Institute, College of Medicine and Public Health, Flinders University, Adelaide, Australia

**Keywords:** Alzheimer’s disease, tau, tau phosphorylation, memory, engram, remote memory, spatiotemporal learning, cued fear conditioning, associative learning

## Abstract

The engram represents the physical trace that encodes a specific memory and enables its recall ^1–4^. Functional failure of the engram is linked to the progressive memory decline in Alzheimer’s disease ^5^. However, it is unknown whether the microtubule-associated protein tau, a central factor in Alzheimer’s ^6,7^, has a direct function in the engram. Here, we demonstrate that tau and encoding-associated tau phosphorylation are critical for robust remote memory engrams. Tau is required specifically during memory formation for remote, yet not proximal recall in memory paradigms in mice. Controlled expression of tau exclusively during memory entrainment is necessary and sufficient to restore remote memory deficits in *tau* knockout mice. Tau is phosphorylated at specific sites during encoding. Gene editing to ablate site-specific phosphorylation at threonine-205 (T205) lowers precision of engram cell recruitment and precludes efficient remote recall. Vector-based engineering of engram cells reveals that T205 phosphorylation of tau is required to engrain memory for recall at remote timepoints. Notably, in the absence of *tau*, memory is recalled from latency by direct optogenetic activation of engram cells at distal time points but not when natural cues are used, revealing an association-specific gatekeeper function of tau during encoding. Our work delineates a physiologic role of site-specific tau phosphorylation at the inception of episodic memory to support an enduring engram and enable efficient remote recall. Thus, encoding-associated phosphorylation of tau is proximal to the elusive substrate of remote memory and may connect to the basis of amnesia in Alzheimer’s disease.

## Main text/ Article

Alzheimer’s disease (AD), the most common form of dementia, is characterised by progressive cognitive decline and loss of existing memory ^8^. Episodic memory deficits occur early in AD and are thought to occur due to ineffective encoding ^9,10^ and impaired memory retrieval ^9^. The engram refers to the physical unit of cognitive information storage in nervous systems ^1–4^. The engram remains in a quiescent state until reactivation, which supports long-term and remote memory and is preferentially reactivated during memory retrieval ^1–4^. For memory to occur, an experience or learning event results in enduring changes in a population or ensemble of neurons recruited to the engram ^2,4,11^. Thus, targeted reactivation of engram cells is sufficient to reinstate memory ^11^. Despite these advances in understanding cellular roles in memory traces, unanswered questions revolve around what defines the basis of the engram’s enduring but latent character, and its underlying molecular nature. Molecules and mechanisms required at memory inception to encode an enduring engram remain unknown, and their loss of function may be directly linked to impairment of long-term memory in AD.

The microtubule-associated protein tau, expressed from the *Mapt* gene, is a neuronal protein that undergoes phosphorylation events in both physiological and pathologic states ^7^. Tau pathology and cognitive decline in AD strongly correlate ^12,13^. Despite tau’s prominence in the context of dementia with amnesia, a direct role of tau in the engram has not been addressed. Here, we show that tau is required for laying down the engram specifically to grant strength to the mnemonic signal and permit efficient retrieval at remote, yet not proximal time points. This function of tau is dependent on its site-specific phosphorylation. This is the first evidence that tau is integral to the molecular nature of the engram.

### Remote memory is impaired in *tau*-deficient mice

Tau is dispensable for memory acquisition and for short- and long-term memory, defined as recall hours and days post-inception, respectively ^14–17^. We therefore asked whether tau is involved in remote memory and assessed standardised paradigms that are commonly employed to study distinct mnemic modes, i.e. aversive and appetitive associative learning or spatiotemporal learning ^18–20^, with recall at very distal timepoints (**Fig. 1a**).

**Figure 1.**
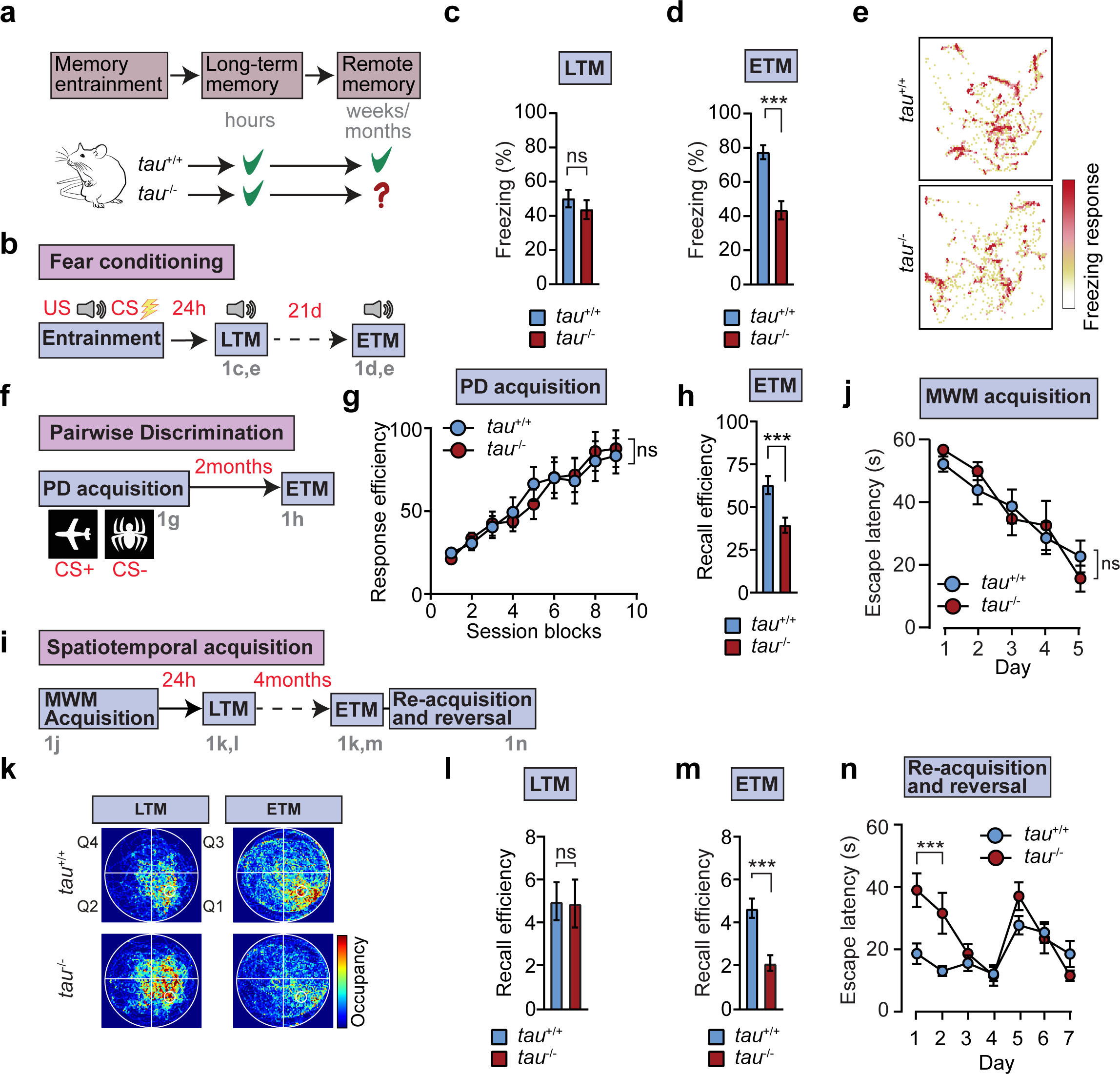
Tau is necessary for remote memory engrams in mice. (**a**) Conceptual schematic – *tau*^+/+^ and *tau*^-/-^ mice show comparable learning and long-term memory. However, recall performance of *tau*-deficient mice at extended time points is unknown. (**b-f**) Impaired remote recall in *tau*^-/-^ mice in cue-dependent associative learning paradigm with aversive enforcer (cued fear conditioning). (n = 18-20) (**b**) Schematic of unconditioned auditory stimulus (US) and conditioned aversive stimulus (CS) pairing and time points of long-term memory (LTM) and extended-term memory (ETM) recall. (**c**) Recall efficiency at long-term time point (24h post-entrainment) (**d**) Recall efficiency at remote timepoint (21 days) (**e**) ETM recall freezing response map. (**f-h**) Impaired remote recall in *tau*^-/-^ mice in appetitive associative learning paradigm (pairwise visual discrimination) (n = 16) (**f**) Example of rewarded (CS+) and non-rewarded response (CS-) pairing, acquisition and recall schedule. (**g**) Acquisition and consolidation efficiency relative to random choice across session blocks in *tau*^+/+^ and *tau*^-/-^ mice. (**h**) Recall efficiency after 2 months. (**i-n**) Impaired remote recall despite initial learning and memory in *tau*^-/-^ mice in a spatial learning paradigm as compared with *tau*^+/+^. (n = 15-20) (**i**) Schematic of spatiotemporal learning and memory paradigm to test long-term (LTM; 24 h) and extended-term (ETM; 4 months) memory recall. Mice are trained to solve water maze, long-term recall is tested after 24 hours, extended-term remote recall after 4 months and mice then re-train in water maze before cue reversal. (**j**) Morris water maze spatial acquisition. (**k**) Occupancy map of *tau*^+/+^ and *tau*^-/-^ mice in LTM and ETM recall trials. (**l**) Recall efficiency at LTM time point (24h post-spatial acquisition) (**m**) Recall efficiency at ETM timepoint (4 months post-spatial acquisition) (**n**) Re-acquisition (d1-4) starting 1 day after ETM recall followed by reversal learning trials (d5-7). Statistical comparisons are performed using ANOVA; ***p<0.001, **p<0.01; ns, not significant. Data are presented as mean ± S.E.M.

We first addressed remote memory in *tau*^+/+^ and *tau*^-/-^ mice in an aversive associative learning paradigm, cued fear conditioning (CFC), which instils a robust engram for cue-response associations measured by freezing response as a proxy of memory recall ^11,19^. After cued conditioning of *tau*^+/+^ and *tau*^-/-^ mice (at 4 months of age), memory was assessed at long-term recall (LTM) on day 1 and at remote or extended-term recall (ETM) 21 days and 42 days post-conditioning (**Fig. 1b**). *Tau^+/+^* mice displayed robust memory at both LTM and ETM recall (**Fig. 1c-e; fig. S1c**). Strikingly, *tau^-/-^* mice exhibited significantly lower ETM responses as compared with *tau*^+/+^ mice (**Fig. 1d-e; fig. S1c**), yet equally efficient recall at LTM (**Fig. 1c, e**), indicating robust memory entrainment and early recall despite a clear deficit in remote recall in the absence of *tau*. Moreover, remote recall deficits were not a consequence of enhanced extinction due to lack of *tau* (**fig. S1d**), precluding active memory degradation and substantiating a role of tau during encoding.

A strong aversive experience, such as the foot shock during conditioning, may engage circuits involved in anxiety-related behaviour ^21^. To control for maladaptive fear responses as cause of reduced post-training freezing in *tau^-/-^* during ETM, locomotor activity and spontaneous freezing behaviour were assessed. Both *tau*^+/+^ and *tau*^-/-^ mice showed similar locomotor activity during habituation (**fig. S1a**), aversive stimulus-induced freezing (**fig. S1b**). Probing anxiety-related behaviour in open-field (**fig. S1e**) and elevated plus maze (**fig. S1f**) corroborated normal anxiety levels in *tau*^-/-^ mice ^17,22^ and confirmed that remote recall deficits are not due to effects on anxiety-related behaviour by *tau* deficiency.

Together these results show tau is required in associative learning with an aversive reinforcer for an enduring engram and memory retrieval at a distal time-point.

Fear conditioning corresponds to associative learning with an aversive stimulus ^19^. To determine whether remote recall attenuation in absence of *tau* applied to memory from appetitive learning, we employed a pairwise discrimination (PD) schedule with remote recall on a touch-screen operant platform, which relies primarily on appetitively-motivated instrumental learning ^20^, in *tau*^+/+^ and *tau*^-/-^ mice (**Fig. 1f**). *Tau*^+/+^ and *tau*^-/-^ mice both reached task acquisition criterion in the PD paradigm with comparable efficiency (**Fig. 1g**). With 24 hours intervals between acquisition trial blocks, long-term memory in the PD task appeared unaffected in *tau*^-/-^ mice. Upon pairwise stimulus exposure 2 months after training, *tau*^-/-^ showed markedly lower recall efficiency as compared with *tau*^+/+^ mice (**Fig. 1h**), which translated into delayed stimulus-award reacquisition (**fig. S2b**). Thus, results in touchscreen-based appetitive learning define a memory deficit in the absence of tau upon remote recall.

To assess remote spatial memory, we optimized a spatio-temporal memory paradigm, Morris water maze (MWM), to include a criterion ensuring robust enduring memory in mice, retrievable at long-term (24hr) and extended-term (>4m) timepoints of memory recall, followed by reacquisition, reversal learning and visual acuity testing (**Fig. 1i**). We compared this protocol of spatial entrainment with non-spatial acquisition controls and thus distinguished chance solution from spatial memory (**fig. S3a-e**).

*Tau*^-/-^ and *tau*^+/+^ mice displayed comparable spatial acquisition to criterion (**Fig. 1j, fig. S1f**) and LTM recall (**Fig. 1k, 1l**), consistent with previous work on navigational learning and spatial long-term memory in *tau*-deficient mice ^7,17,23,24^. Swim path analysis verified that *tau*^-/-^ mice did not employ compensatory mechanisms during spatial acquisition (**fig. S3f**). Strikingly, genetic deletion of tau (*tau*^-/-^) significantly impaired remote recall efficiency as compared with *tau^+/+^* mice (**Fig. 1m**), which was corroborated by averaged maze occupancy (**Fig. 1k**) and annulus crossing analysis of probe trials at LTM and ETM (**fig. S3g**).

Overall, these results demonstrate that enduring spatial memory requires tau for efficient retrieval at remote timepoints. Significantly reduced extended-term recall efficiency in aversive associative, appetitive, and spatial learning paradigms indicate that different mnemic processes all require tau for adequate remote recall.

### Tau expression during encoding is necessary and sufficient for robust remote recall

The remote recall deficit in *tau*^-/-^ was not a result of altered extinction as fear response to extinction trials at different time points post-conditioning were comparable between *tau*^+/+^ and *tau*^-/-^ (**fig. S1d**), suggesting that tau was important during initial entrainment of remote fear memory. We therefore hypothesized that re-introducing tau during encoding may restore remote memory retrieval. To investigate this concept, we used an adeno-associated virus (AAV)-based vector system for neuron-specific expression of tau or enhanced green fluorescent protein (eGFP) solely during memory acquisition in spatial and associative learning paradigms (MWM and CFC, respectively) (**Fig. 2a**). Notably, both spatial memory and remote fear memory are hippocampal-dependent ^25–27^. The AAV system is under control of an improved tetracycline transactivator (d2tTA), expressed from a neuron-specific *synapsin-1* (*syn1*) promoter, for inducible expression and is delivered stereotactically to the murine hippocampus as a single viral vector (**Fig. 2a**) ^28^. Transgene expression is suppressed by providing doxycycline (DOX)-containing feed, preventing d2tTA from binding to the TRE element, and induced by relieving the DOX block with regular DOX-negative chow (**Fig. 2a**).

**Figure 2.**
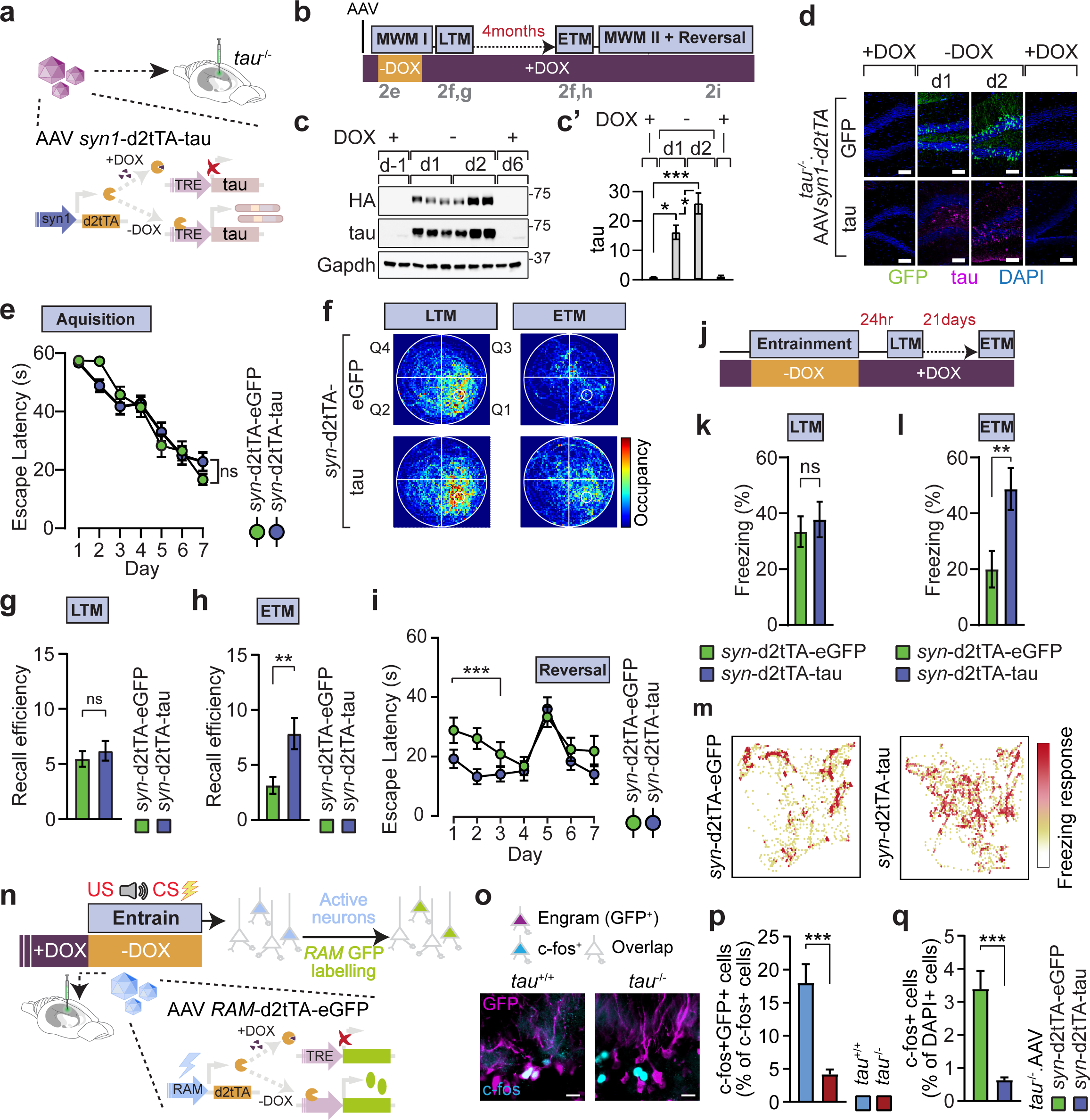
Experience-restricted neuronal tau expression is sufficient and required for remote recall. (a) Schematic of controlled neuronal tau expression in *tau*^-/-^ mice. Neuron-specific minimal promoter (*syn1*) under control of doxycycline, which expresses tau or eGFP in the absence of doxycycline (-DOX). In the presence of DOX (+DOX) tetracycline transactivator (d2tTA) binds to DOX and prevents expression. Construct packaged in a single adeno-associated virus (AAV) vector are bilaterally delivered into the hippocampus the of *tau*^-/-^ mice. (b) Schematic of MWM schedule with long-term and remote recall Schematic representation of controlled tau expression during Morris water maze (MWM). During memory acquisition (MWMI) tau expression is instated in the absence of DOX (-DOX). Following acquisition, DOX is administered (+DOX) and tau expression is prevented. Extended-term memory (ETM) is measured 4-months post-acquisition, followed by re-learning (MWMII) and long-term memory (LTM) testing before reversal learning and visual acuity testing. (c) Representative immunoblot of hippocampal *tau*^-/-^ lysates showing tight control of inducible tau expression from neuron-specific d2tTA AAV system (n=2-3). Immunoblots are probed for exogenous HA-tagged tau, total tau, and Gapdh as loading control. In the presence of DOX (+DOX) no tau expression is detected on day -1 (d-1). On day 1 (d1) and 2 (d2) following DOX removal (-DOX), tau expression is detected. Reinstatement of DOX (d6) results in suppression of tau expression. (**c’**) Quantification of immunoblot in d (n=4-6). (d) Immunofluorescence analysis confirming eGFP or tau expression following DOX removal in hippocampal sections from *syn*-d2tTA-tauWT or control *syn*-d2tTA-eGFP (n=4). Scale bar, 100µM. (e) Escape latencies during acquisition phase. Graph represents escape latency (s) of 3-month *tau*^-/-^ mice with indicated AAV treatment (*syn*-d2tTA-tauWT or control *syn*-d2tTA-eGFP) (n=22-23 mice per AAV). No difference in escape latency (s) is observed across both groups. (f) Occupancy map of *tau*^-/-^ mice with indicated AAV treatment in LTM and ETM recall trials (n=20-24 mice per AAV). (g) Long-term memory (LTM) probe trial (n=22-23 mice per AAV). Graph represents recall efficiency (s) during LTM test assessed by occupancy in water maze target quadrant. No differences observed between both groups for target quadrant occupancy. (h) Extended-term memory (ETM) probe trial (n=22-23 mice per AAV). Tau^-^injected mice display significantly longer occupancy in target quadrant compared to eGFP control mice.. (i) Re-acquisition (d1-4) 4 months after memory recall in e followed by reversal control trials (d5-7) (n=20-24 mice per AAV). (**j-m**) Cued fear conditioning combined with neuronal d2tTA system-mediated hippocampal expression of eGFP or tau. *Tau*^-/-^ mice with indicated AAV treatment (*syn*-d2tTA-tauWT or control *syn*-d2tTA-eGFP) were habituated on DOX (+DOX) 24hrs before entrainment (-DOX). Following entrainment mice were placed on DOX (+DOX) for LTM (24hrs post) and ETM (21days post entrainment) test. (k) LTM test for *tau*^-/-^ mice with indicated induced expression during CFC entrainment (n=21-22). LTM recall (24 h) is comparable for induced eGFP and tau expression during encoding. (l) ETM test (21 days) for *tau*^-/-^ mice with indicated induced expression during CFC entrainment (n=21-22). ETM recall is markedly better after tau expression during encoding. (**m**) ETM recall freezing response map. (n) Schematic for hippocampal engram labelling using robust activity marking (*RAM*) promoter-driven eGFP expression during encoding of fear association. (o) Immunofluorescence for activity marker c-fos in hippocampus of upon tau and eGFP expression with AAV *syn*-d2tTA system during entrainment in CFC (or MWM) in *tau*^+/+^ and *tau*^-/-^. (p) Overlap of engram (eGFP^+^) with c-fos^+^ cells in *tau*^+/+^ and *tau*^-/-^ during fear memory encoding. (n=6-7) (q) Number of c-fos^+^ cells in hippocampus of *tau*^-/-^ mice expressing eGFP or tau during fear memory encoding (*tau*^-/-^.AAV syn-d2tTA-eGFP/tau). (n=6-9) Statistical comparisons are performed using ANOVA; ***p<0.001, **p<0.01, *p<0.05; ns, not significant. Data are presented as mean ± S.E.M.

We employed this system in *tau*^-/-^ mice to restrict neuron-specific hippocampal expression of tau to spatial acquisition in the MWM followed by DOX-mediated suppression of tau for 4 months and during subsequent ETM recall and spatial reacquisition (**Fig. 2b**). Immunoblot of hippocampal lysates confirmed that expression increased over acquisition days 1 and 2 following DOX removal, while expression was completely suppressed once DOX was reinstated (**Fig. 2c**). Immunofluorescence confirmed restricted expression to the hippocampus, rapid induction from 12h post-DOX removal onward, and suppression of tau and eGFP by reinstatement of DOX feed (**Fig. 2d**, **fig. S4**). Both *tau*^-/-^ mice injected with AAV *syn1*-d2tTA-tau or AAV *syn1*-d2tTA-eGFP and induced expression during the spatial acquisition phase showed comparable learning (**Fig. 2e**) and LTM recall (**Fig. 2f, 2g, fig. S5b**). Strikingly, ETM recall (**Fig. 2f, h, fig. S5c**) and reacquisition (**Fig. 2i**; days 1-3) were significantly better in *tau*^-/-^ mice expressing tau during initial spatial acquisition as compared with mice expressing eGFP in the same time window 4 months prior. Notably, temporally learning-restricted tau expression did not modulate reversal learning compared to controls (**Fig. 2i**; days 5-7). Thus, neuronal tau expression restricted to encoding is necessary and sufficient to support robust recall at extended term timepoints, while dispensable for more proximal memory. Moreover, these results indicate that tau is not required to prevent memory degradation over time because of efficient memory recall at ETM timepoints (**Fig. 2h**; **fig. S5c**) despite lack of tau expression during the latency (when DOX-containing feed is supplied; **Fig. 2c-d; fig. S4**). Instead, tau is required during the initial encoding to enable efficient ETM recall.

To examine if learning-restricted tau expression facilitates remote memory recall of associative learning experiences, LTM (24hr) and ETM (21d and 42d) in CFC was quantified in *tau*^-/-^ mice injected with AAVs *syn1*-d2tTA-tau or -eGFP. Expression was permitted solely during entrainment and suppressed during habituation, latency, and recalls (**Fig. 2j**). Both groups displayed similar basal activity levels during habituation, training-induced freezing during entrainment (**fig. S6a-b**), and LTM recall of tone-shock association (**Fig. 2k**). Strikingly, eGFP-injected mice were impaired for ETM (**Fig. 2l, m; fig. S6c**), which suggests neuronal tau is required for entrainment and recall of remote associative memories.

Taken together, results from spatial and associative learning paradigms with temporally controlled reinstatement of neuronal tau expression demonstrate a critical function of tau for memory recall at distal time points, which is distinctly required during the learning experience.

Temporally controlled tau expression may induce changes in local network activity, which is involved in mnemic processes ^29^, and thus facilitate encoding and recall. To assess hippocampal circuit activity in mice with temporally controlled tau expression, we employed telemetric EEG recording (**fig. S7**). No differences in EEG power spectrograms and cross-frequency coupling, a proxy of network connectivity supporting hippocampal memory function ^30,31^, were observed in presence (-DOX) or absence (+DOX) of tau or eGFP in *tau*^-/-^ mice. EEG results suggest comparable local network activity in mice expressing tau or eGFP during learning experience and beyond. This is consistent with hippocampal network and electrophysiologic normality ^16,32,33^, and normal hippocampal learning, short and long-term memory (up to 24h) in *tau*^-/-^ mice ^14,16^.

We therefore investigated effects on a cellular level. Cells active during encoding form part of the engram i.e., ensembles responsible for learning-induced changes ^2,11^. Neurons that are rendered more excitable during fear conditioning have a greater likelihood of being allocated to the memory ensemble ^34^. However, sparsity of population activity is critical for discrimination of sensory input patterns and precision of association encoding ^35–37^. Neuronal activity induces immediate early gene (IEG) expression, which serves as a molecular signature for marked activation ^38^. We examined to what extent neurons were represented in the fear memory engram in *tau*^+/+^ and *tau*^-/-^ hippocampus. To visualise the memory trace, we labelled CFC engram with eGFP expression under control of activity-dependent promoter ^39^ and stained for IEG marker c-fos ^38,40^ (**Fig 2n**). We assessed abundance and overlap between c-fos^+^ and engram-associated (eGFP^+^) neurons (**Fig. 2o; fig S8a**), which showed that while *tau*^-/-^ displayed more c-fos^+^ cells (**fig S8b**), proportionally fewer c-fos^+^ cells were allocated to the engram (**Fig. 2p**; **fig S8c**). Consistently, *tau*^-/-^ mice expressing tau from AAV *syn1*-d2tTA-tau during entrainment had fewer c-fos^+^ cells as compared with *tau*^-/-^ injected with AAV *syn1*-d2tTA-eGFP (**Fig. 2q**). These results suggest that tau limits activation during entrainment in non-engram cells and supports sparse hippocampal cell activation, which enhances precision and effective mnemonic signal for remote memory ^36,37^.

Taken together, experiments with experience-restricted tau expression show that the critical function of tau for remote memory recall (*1*) is mediated by hippocampal tau solely during the entrainment and independent of tau during memory latency; (*2*) is independent of local network activity changes; and (*3*) correlates with precision of encoding-specific recruitment of cells to engram ensembles. This demonstrates that tau during memory encoding, yet not during the subsequent latency phase, is required and sufficient for robust remote recall of the engram.

### Site-specific T205 tau phosphorylation promotes remote memory

Tau occurs in phosphorylated forms in physiologic and pathologic states ^6,41,42^, and phosphorylation of tau modulates learning in rodent models of memory impairment ^16,43,44^. A physiologic role of tau phosphorylation specifically in the engram is however unknown. We asked whether tau phosphorylation was involved during learning experience. To address this, specific sites of tau phosphorylation were probed in tau expressed during spatial acquisition in *tau*^-/-^.AVV *syn1*-d2tTA-tau mice by immunoblot with antibodies of confirmed phospho-epitope specificity ^44^ (**Fig. 3a**). Phosphorylation of tau was readily detectable at serine-202 (S202) and threonine-205 (T205) and to a lesser extent at T181 in the proline-rich region of tau, while modification at C-terminal sites S396, S404 and others were not detectable (**Fig. 3b-b’**; **fig. S3-9**). T205 phosphorylation was readily detectable in tau^+^ hippocampal cells of *tau*^-/-^.AVV *syn1*-d2tTA-tau mice at day 1 of spatial acquisition (**Fig. 3b**). Thus, specific phosphorylation is induced in tau expressed in hippocampal neurons during physiologic learning.

**Figure 3.**
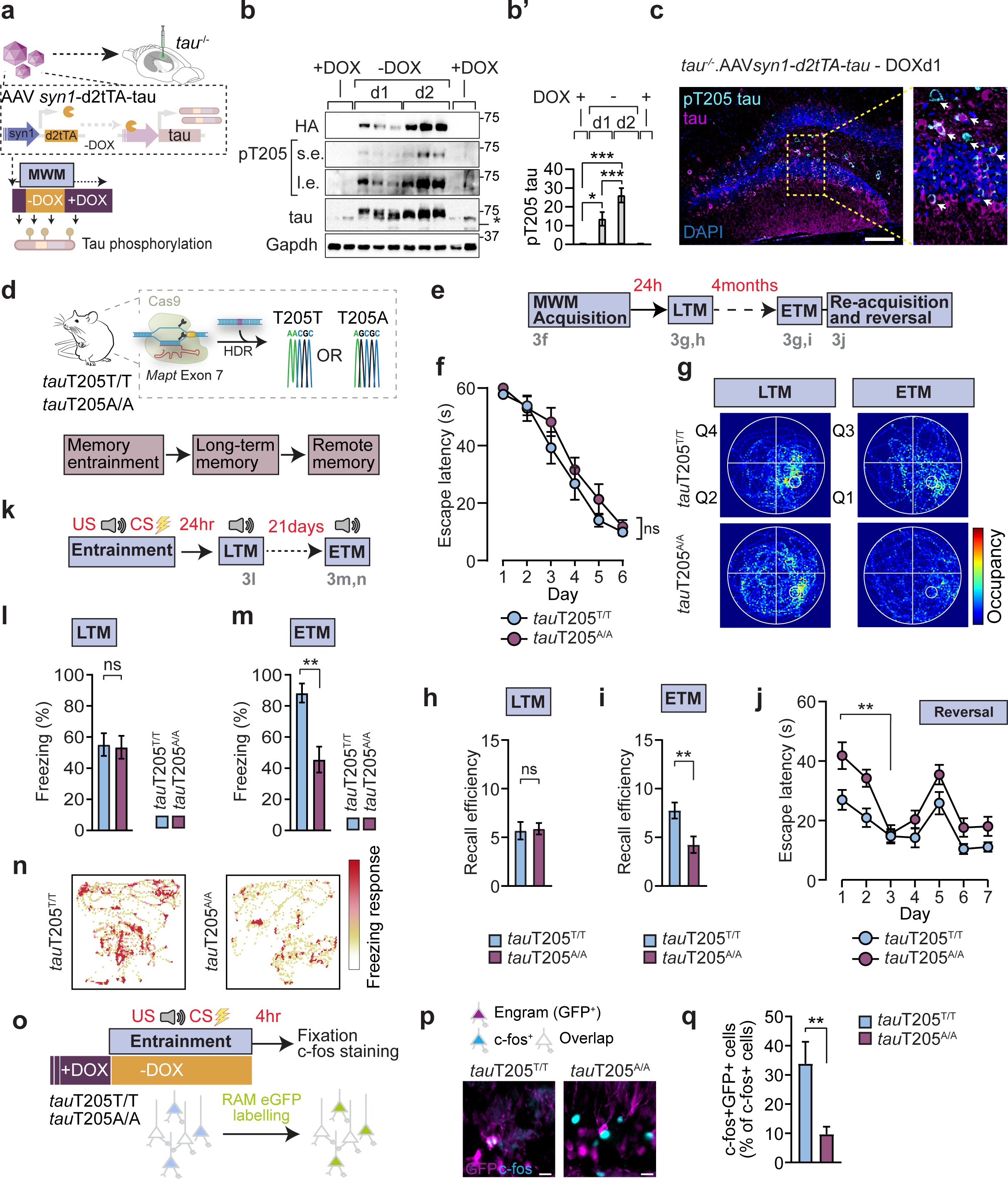
Site-specific T205 tau phosphorylation mediates remote memory. (**a**) Schematic of tau phosphorylation analysis at different timepoints of controlled neuronal expression and memory entrainment schedule. (**b – b’**) Representative immunoblot of specific phospho-Threonine205 (pT205) epitope on tau expressed in *tau*^-/-^ hippocampus during spatial memory acquisition using neuron-specific DOX-dependent d2tTA expression system (n=2-3). d1-2, day 1-2 off DOX. HA, haemagglutinin tagged exogenous tau. Gapdh, loading control. s.e., short exposure; l.e., long exposure. (**b’**) Quantification of immunoblot in a for pT205 tau normalised to total tau (n=4-6). (**c**) Immunofluorescence labelling for pT205 tau, tau and GFP of hippocampal sections from *tau*^-/-^ mice injected with AAV *syn1*-d2tTA-tau or -eGFP at day 1 (d1) after removal of DOX-containing diet (-DOX) and start of spatial acquisition. DAPI, nuclear labelling. Scale bar, 200 µm. (**d**) Schematic of memory entrainment, long-term and remote recall experiments in mice with CRISPR-engineered *tau*T205A allele. HDR, homology directed repair (**e-j**) Impaired recall at remote timepoint of spatial acquisition memory in *tau*T205A/A mice as compared with *tau*T205T/T mice (n=18-20). (**e**) Spatial acquisition and memory recall schedule in Morris water maze (MWM) paradigm. (**f**) MWM spatial acquisition (**g**) Occupancy map of *tau*T205T/T and *tau*T205A/A mice in LTM and ETM recall trials (**h**) Recall efficiency at LTM time point (24h post-spatial acquisition) (**i**) Recall efficiency at ETM timepoint (4 months post-spatial acquisition) (**j**) Re-acquisition (d1-4) starting 1 day after ETM recall followed by reversal learning trials (d5-7). (**k-n**) Impaired remote fear memory recall in *tau*T205A/A mice (n = 15-16) (**k**) Schematic of cued fear conditioning schedule (unconditioned auditory stimulus (US) and conditioned aversive stimulus (CS) pairing) including time points of LTM and ETM recall. (**l**) LTM recall efficiency (24h post-entrainment) of *tau*T205T/T and *tau*T205A/A mice (**m**) ETM remote recall efficiency (21 days) of *tau*T205T/T and *tau*T205A/A mice (**n**) ETM recall freezing response map. (**o**) Schematic for hippocampal engram labelling using robust activity marker (RAM) promoter-driven eGFP expression in *tau*T205T/T and *tau*T205A/A mice 4h post-entrainment of CFC association. (**p**) Co-labelling of hippocampal engram cells (eGFP^+^) with activity marker c-fos in *tau*T205T/T and *tau*T205A/A mice during fear memory encoding. (n=5). Scale bar, 20 µm. (**q**) Overlap of fear memory engram (GFP^+^) with c-fos^+^ cells from *tau*T205T/T and *tau*T205A/A mice (n=5-6). Statistical comparisons are performed using ANOVA; ***p<0.001, **p<0.01, *p<0.05; ns, not significant. Data are presented as mean ± S.E.M.

Site-specific T205 phosphorylation of tau is a target of post-synaptic kinase p38γ ^45^, albeit its physiologic relevance is currently unknown. Prominence of phospho-T205 in learning-associated tau led us to investigate the role of this site in learning and recall at long-term and remote timepoints, using mice with CRISPR-edited T205 codon in the endogenous *Mapt* gene, in which phosphorylation is precluded by a T → Alanine (A) codon exchange (*tau*T205A)^45^ (**Fig. 3d**).

*Tau*T205T/T and *tau*T205A/A mice were subjected to a MWM schedule (**Fig. 3e**) and showed equal spatial learning (**Fig. 3f**) and initial memory (LTM) (**Fig. 3g, h**), as in prior studies ^45^. Strikingly, ablation of T205 impaired remote memory retrieval in ETM trials (**Fig. 3g, i; fig. S9b**), indicating mice have diminished remote memory for the spatiotemporal task compared to wildtype *tau*T205T/T mice. Ablation of T205 did not impact on spatial reacquisition or reversal learning (**Fig. 3j**), demonstrating T205 is dispensable for spatial learning and memory, with the exception of extended-term recall. Mice lacking T205 tau kinase *p38γ* (*p38γ*^-/-^) demonstrated a similar phenotype for remote recall, while initial spatial acquisition and LTM recall were comparable to *p38γ*^+/+^ controls (**Fig. S10**), the latter being consistent with previous results showing no requirement of p38γ for initial memory acquisition ^16^.

We next utilized CFC (**Fig. 3k**) and confirmed preventing T205 phosphorylation does not impact on LTM (**Fig. 3l; fig. S12a-b**). However, much like in *tau*^-/-^ mice, ablation of T205 impaired freezing behaviour during ETM (**Fig. 3m, n**) trials. Extinction of fear memory after entrainment was comparable between *tau*T205T/T and *tau*T205A/A mice, precluding a role of T205 tau phosphorylation in active uncoupling of the fear response (**fig. S12c**). To ensure that ablating T205 had no impact on generalised anxiety-related behaviour, we performed elevated plus maze tests (**fig. S12d**). Overall, these results confirm T205 is an important regulatory site for remote memory and underlies the tau-dependent mechanism that gates formation of memory for distal retrieval. This is the first evidence of a physiologic requirement of an individual tau phosphorylation site for remote memory.

To examine whether tau phosphorylation at T205 was involved in the neuronal ensemble associated with the engram, we labelled the fear memory engram with eGFP and stained for IEG c-fos (**Fig 3o**). While c-fos^+^ cells were more abundant in tauT205A/A (**fig. S13a-b**), fewer c-fos^+^ neurons were recruited to the labelled engram (**Fig. 3p-q; fig. S13c**), indicating that T205 phosphorylation of tau is involved in the active cell ensemble associated with the engram at the time of encoding.

### Encoding-specific tau phosphorylation in the engram supports remote memory

For more direct evidence of a role of active ensemble-linked tau phosphorylation for remote memory, we developed a vector to directly query tau in engram cells for memory encoding, long- and extended-term recall. To target hippocampal cells active during encoding, which form part of the engram, we devised an AAV expression system that combines a Robust Activity Marking (*RAM*) promoter, an activity-dependent gene expression element of enhanced sensitivity and selectivity, with the d2tTA-TRE system for tight temporal control of engram-specific targeting ^39,46^ (**Fig. 4a**). We modified this system to express wild-type tau (tauWT) or phospho-T205-defective tauT205A in memory ensembles or to label engram cells with a fluorescent marker (eGFP)(**Fig. 4a**). Delivery of AAV particles to the hippocampus of *tau*^-/-^ mice and removal of DOX-containing diet during memory entrainment in MWM or CFC labelled hippocampal engram cells with either eGFP (**Fig. 4b**) or expressed tauWT or tauT205A (**Fig. 4b, fig. S14a**). Engram-specific tauWT showed T205 phosphorylation, which was absent in the tauT205A variant-expressing engram (**Fig. 4b, c; fig. S14b**). Engram expression of tauWT or tauT205A had no effect on local network activity in EEG recordings in *tau*^-/-^ mice (**fig. S15**). However, engram-specific expression of tauWT but not tauT205A limited hippocampal c-fos^+^ reactivity (**Fig. 4d, 4d’**), corroborating a limiting effect of tau in engram cells on proximal neuronal activation to enhance precision of information encoding.

**Figure 4.**
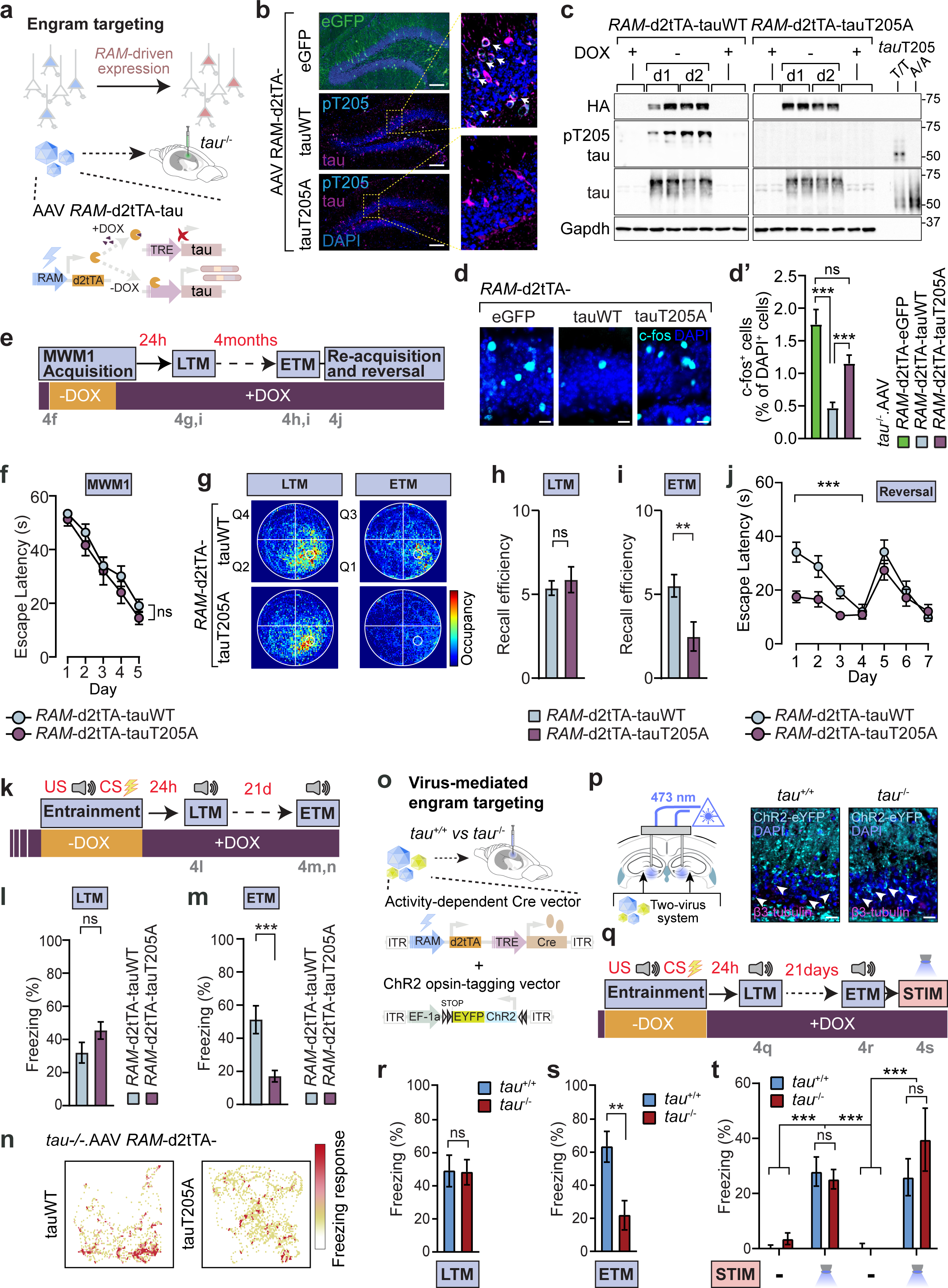
Remote memory engrams rely on encoding-associated site-specific tau phosphorylation. (a) System for engram-specific tau expression based on *RAM* promoter and doxycycline (DOX)-controlled d2tTA-TRE system in adeno associated virus (AAV) vector delivered to the hippocampus of *tau*-deficient mice (*tau*^-/-^). In the absence of DOX, expression of wildtype tau (tauWT) or tauT205A variant are induced specifically in cells active during entrainment. (b) Representative images of hippocampus regions showing tauWT, tauT205A and eGFP (green) labelled active cells (d1 -DOX) with immunofluorescence for pT205, tau and GFP. DAPI, nuclear label. Scale bar, 100 µm. (c) Immunoblot of hippocampal lysates from tau-/-mice injected with AAV *RAM*-d2tTA-tauWT or -tauT205A in the presence (+DOX) or absence (-DOX) of doxycycline in diet for indicated days during spatial acquisition in water maze (n=2). (d) Immunolabelling of hippocampal cells in *tau*^-/-^ mice with *RAM*-driven expression of eGFP, tauWT or tauT205A during fear memory encoding cells with activity marker c-fos. DAPI, nuclei; scale bar, 20 µm (**d’**) Percent of c-fos^+^ cells in (d) (n=6) (**e-j**) Engram-specific expression of tauWT or tauT205A during spatial acquisition in MWM and impact on LTM and ETM recall efficiency. (n = 14-15) (**e**) Experimental outline and engram labelling strategy. *Tau*^-/-^ mice are injected in the hippocampus with AAV *RAM*-d2tTA-tauWT or -tauT205A whilst on DOX feed. During acquisition (MWM1) mice are placed on chow diet (-DOX) to enable engram labelling. Following, acquisition mice are placed on DOX (+DOX) and kept on DOX continuously. LTM is tested 24h, ETM 4-months post-acquisition. Mice undergo re-acquisition in the MWM. In the final stage cognitive flexibility is tested (reversal learning). (**f**) Normal spatial acquisition in *tau*^-/-^ mice with engram-specific expression of tau and tauT205A. Graph represents escape latency (s) of tau^-/-^ with indicated AAV (AAV *RAM*-d2tTA-tauWT or AAV *RAM*-d2tTA-tauT205A (n =14-15 per AAV). No differences between both AAV groups. (**g**) Occupancy maps for LTM and ETM probe trials for indicated experimental groups (**h**) LTM recall in probe trial 24 hours post-spatial acquisition in *tau*^-/-^ mice with indicated engram-specific expression. Both groups show comparable LTM recall. (**i**) ETM recall in probe trial 4 months post-spatial acquisition in *tau*^-/-^ mice with indicated engram-specific expression. Engram-specific expression of tauT205A variant during spatial acquisition is insufficient to mediate ETM recall. (**j**) Re-acquisition and reversal learning. Graph represents escape latency (s) for spatial re-acquisition (days 1-4) and reversal learning (days 5-7). (**k-n**) Engram-specific expression of tauWT, yet not tauT205A variant during conditioning is sufficient to mediate ETM recall in *tau*^-/-^ mice (**k**) Cued fear conditioning and engram ensemble expression schedule for entrainment of associative memory. *Tau*^-/-^ mice with indicated AAV treatment (*RAM*-d2tTa-tauWT or *RAM*-d2tTa-tauT205A) were habituated on DOX (+DOX) 24hrs before entrainment (-DOX), which induces expression of transgenes in engram ensembles. Following entrainment mice were placed on DOX (+DOX) for LTM (24hrs post-entrainment) and ETM (21days) testing. (**l**) LTM recall for associative fear memory in *tau*^-/-^ mice with engram tauWT or tauT205A expression during entrainment (n =16-18). (**m**) ETM fear memory recall in *tau*^-/-^ mice with engram tauWT or tauT205A expression during entrainment. (n =16-18). (**n**) ETM recall freezing response map. (**o-t**) Expression of fear memory at remote time point by direct optogenetic activation of engram ensemble but not by conditioned stimulus (CS) presentation in the absence of *tau*. (**o**) Dual AAV engram targeting strategy with blue light-sensitive opsin (ChR2^H134Y^-YFP) in the hippocampus of *tau*^+/+^ and *tau*^-/-^ mice (**p**) Opsin expression in ensembles active during entrainment. Immunolabelling of ChR2-YFP, β3-tubulin and nuclei (DAPI) in hippocampus upon RAM-driven cre-dependent expression of ChR2-YFP. Arrows indicate ChR2-YFP-expressing dentate gyrus cells. Scale bar, 20µm (**q**) DOX block removal during CFC entrainment in context A. LTM and ETM recall in context A with CS at 24h and 21d, respectively. Blue light-induced reactivation at day 22 in context B without CS (STIM). (**r**) LTM, (**s**) ETM recall and (**t**) light-induced response in indicated mice and context (n=8-9). Statistical comparisons are performed using ANOVA; ***p<0.001, **p<0.01, *p<0.05; ns, not significant. Data are presented as mean ± S.E.M.

We next induced memory cell-specific tau variant expression to study the role of tau T205 phosphorylation specifically in the engram generated in the MWM spatiotemporal paradigm and assess LTM and ETM recall (**Fig. 4e**). We induced tauWT or phospho-defective tauT205A expression during memory acquisition (-DOX) and then tested LTM (+DOX) and ETM (+DOX). Both groups of mice showed normal learning (**Fig. 4f**), LTM recall (**Fig. 4g, h** **fig. S16b**), and reversal learning (**Fig. 4j**). However, tauT205A expression in the forming engram was insufficient to support efficient ETM recall as compared with mice with tauWT in engram cells during spatial acquisition, and thus, impaired remote memory (**Fig. 4g, i; fig. S16c**) and delayed spatial reacquisition (**Fig. 4j**). This data demonstrates control of robust encoding of remote spatial memory by site-specific T205 phosphorylation in engram cells.

We next asked whether T205 phosphorylation in the engram governs memory in associative learning. To restrict activity-dependent tau expression to the engram ensemble solely during memory formation, mice remained on DOX feed prior to CFC entrainment and returned to it afterwards (**Fig. 4k, fig. S17a-c**). Engram-specific tauT205A-expression in *tau*^-/-^ mice resulted in normal learning and LTM compared to tauWT expression in engram cells (**Fig. 4l**). Strikingly, lack of T205 phosphorylation in engram-specific tau during entrainment was accompanied by lower remote memory retrieval while tauWT expression supported sufficient ETM recall (**Fig. 4m, n**). Thus, engram-bearing cells of fear memory require T205 phosphorylation to encode a stable and enduring memory, and introducing a form of tau lacking the T205 site is insufficient to instil memory for robust remote recall.

Tau may be involved at memory inception to induce changes in the engram ensemble related to neuronal activation and subsequent expression of memory. Alternatively, tau may be linked to memory substrates independent of neuronal activation. We therefore asked whether tau is required for memory expression at remote time points upon direct neuronal activation of the engram ensemble or whether tau specifically makes physiologic recall induced by natural cues permissive. To test these possibilities, we turned to optogenetic stimulation of engram ensembles at remote time point in presence or absence of *tau*. To make ensembles susceptible to stimulation, the engram was labelled specifically during entrainment in CFC (+CS in context A) with ChR2-YFP by dual hippocampal injection of a *RAM*-driven cre recombinase and a double floxed ChR2-YFP AAV into *tau*^+/+^ and *tau*^-/-^ mice combined with a DOX feed schedule to target cells within the fear engram (**Fig. 4o-q; fig S18**). Mice were then probed for memory (+CS in context A) at LTM and ETM timepoints, followed by light-induced reactivation in absence of stimulus and in an unfamiliar context (-CS in context B) (**Fig. 4q**). Again, ETM recall was impaired in *tau*^-/-^ mice, while LTM recall was comparable to *tau*^+/+^ (**Fig. 4r-s**). Strikingly, activation of the opsin-labelled engram at day 22 post-entrainment was sufficient for memory expression (**Fig. 4t**). Thus, engram retrieval from latency at remote time points is amenable to direct ensemble activation in the absence of *tau*, indicating a gate keeper function of tau that renders natural stimulus-induced recall permissible and distinguishes remote memory recall from purely neuronal activation of the ensemble of the original inception.

In summary, our study describes a role of tau during memory formation that is critical for retrieval, specifically at distal time points. Presence of tau restricted to learning experiences is sufficient to promote remote memory recall in multiple forms of learning. Tau supports learning involving synaptic plasticity in the short and long term ^43,47^. Yet, previous memory studies with *tau*-deficient mice focussed on working, short-term or long-term memory, showing no or minor effects of absence of *tau* ^7,14,16,17,24,47,48^, which vary with gene targeting strategy and genetic background ^49^. Involvement of physiologic tau for remote memory function remained elusive.

A role of tau that enables remote memory, yet not long-term memory is consistent with different underlying mechanisms of persistent mnemonic signals over months or years than over hours or days^3^. We show that tau’s function in this context is inherently linked to site-specific tau phosphorylation. Engram tau is phosphorylated, and encoding-specific tau phosphorylation at T205 is critical for precise neuronal allocation to the engram and limiting proximal neuronal activity, in accordance with the need for sparsity and competitive inhibitory control in the engram ^36,37^. Moreover, the concept of synaptic tagging of memory traces proposes an early distinctive molecular signal at encoding that influences, through as of yet unknown mechanisms, hippocampal-coordinated storage and is critical for formation and temporal persistence of remote memories ^3^. Site-specific tau phosphorylation, based on our findings, may be part of this mechanistic link. Notably, AD pathology presents with a high load of tau phosphorylated at the AT8 epitope, which includes T205 ^50,51^, potentially interfering with the physiologic biochemical tau signature in the engram at the outset of lasting and robust memory.

Notably, the tau protein is dispensable during latency. Tau has a significant impact on remote recall in *tau*-deficient mice when expression is restricted to experience learning despite complete suppression during latency. This indicates that tau, while not the primary resin of memory in an engram ensemble, governs lasting memories at encoding in a molecular event proximal to the substrate of memory and is sufficient and required to induce its enduring nature.

Episodic and visuospatial memory are affected early in progressive tauopathies such as Alzheimer’s disease or dementia with Lewy bodies ^9,52^, and amnesia in dementia tauopathies is presently irreversible ^53^. The current tenet holds that tau pathology impairs cellular and circuit functions in limbic regions subserving episodic memory ^6,50,54^, leading to an early degradation of episodic recollection ^9,55^ and visuospatial abilities ^56^. Direct optogenetic activation of engram ensemble facilitates remote recall in absence of tau, indicating that tau – at the outset of a learned experience – makes encoded memory permissible to natural remote recall specifically. This role of tau in the engram reconciles concepts of impaired encoding ^9,10^ and impaired retrieval ^5^ of memories in AD. Thus, the importance of encoding-associated phosphorylation of tau in the engram, newly identified in this study, is critical to advance our understanding of remote memory and the basis of memory loss in dementia.

## Author contributions

A.I. developed the concept of this study and overall research strategy.

K.S. and A.I. designed the study and all experiments.

K.S. and A.I. wrote the manuscript with contributions from E.P., R.K., and Y.L.

K.S., R.K., and A.I. performed all animal experiments, tissue isolation, histology, imaging, and image analysis.

K.S. and A.I. performed memory test analysis.

K.S., E.P., Y.L., and A.I. generated all viral vectors and plasmid constructs used in this study.

## Ethics declarations

A.I. is a co-inventor on a patent application related to targeting p38γ and Thr-205 tau in Alzheimer’s and other neurodegenerative diseases (Australian patent number APA#2016900764).

## Acknowledgements

We thank Dr Yee Lian Chew, Dr Fabien Delerue, and Prof Lars Ittner for discussions on experimental design. Holly I Ahel, Alexander M Volkerling and Prita R Asih for technical assistance. Prof Karin Nordstrom and Dr Yee Lian Chew for critical reading of the manuscript. Flinders Microscopy (Dr Nicholas Eyre, Pat Vilimas) for assistance with histology and microscopy. Technical staff of the Flinders Biomedical Engineering and the College of Medicine and Public Health Animal Facility of Flinders University.

## Funding

This work was supported by funding to A.I. from the National Health and Medical Research Council (grant nos. 1143978 and 1176628), the Australian Research Council (grant nos. DP200102396, and DP220101900), the Dementia Australia Research Foundation, the Flinders Foundation, and Flinders University and by funding to K.S. from the BrightFocus Foundation (grant no. A2022022F to K.S.) and Dementia Australia Research Foundation. This work was supported by funding to E.P. from Flinders Foundation. A.I. is a National Health and Medical Research Council Emerging Leadership 2 fellow (grant no. 1176628). K.S. is a Scientia Professor Henry Brodaty Post-doctoral Fellow of the Dementia Australia Research Foundation.

## Methods

### Mice

All animal experimentation and husbandry were approved by the Animal Welfare Committee of Flinders University and followed Institutional Animal Care and Use Committee guidelines. Mice with targeted alleles for *p38*γ (*Mapk12*) [*p38*γ^−^; B6.129S4(Cg)-*Mapk12^tm^*^1^*^.2Lex^*]^57^, *tau*-deficient mice [tauEx1; B6.129S4(Cg)-Mapt]^22^, and *tau*T205A mice with the *Mapt* gene targeted by CRISPR-Cas9 to exchange the codon corresponding to T194 (i.e., T205 in human tau)^45^ were previously described. All mouse strains were maintained on a C57Bl/6 background. Mice were housed in individually vented cage systems (Tecniplast, Italy) with access to food and water ad libitum (Gordon Specialty Feeds, standard rodent chow). Lighting was adjusted to a 12-hour circadian day-night cycle. Regular health checks were carried out by animal facility staff and researchers, and only animals with normal health parameters (body weight and coat condition) were included in experimentation. All testing and analyses were done by researchers blinded to genotypes and treatments. Mice were genotyped by polymerase chain reaction (PCR) using isopropanol-precipitated genomic DNA as template and OneTaq DNA polymerase (New England Biolabs, (NEB), #M0480) as described previously ^44^. *Tau*T205A mice were genotyped using isopropanol-precipitated DNA from ear biopsies as template for tetra-primer amplification-refractory mutation system (4ARMS) PCR ^45^. Oligonucleotide primers (desalted; Sigma) for genotyping targeted alleles and transgenes by PCR are described in Supplementary Table 2.

### Memory and behaviour testing in mice

#### Fear conditioning

Fear conditioning protocols were based on previously published studies ^11^ and performed on a system with automated load cell detection, shock grid floor and video monitoring (PanLab, Spain). The chamber floor consisted of metal parallel grid bars connected to the load cell coupler and shock generator. The conditioning chamber was placed inside a custom-built sound attenuating box with inbuilt ventilating fan. A camera was custom built into the ceiling of the fear chamber for visualisation and analysis. The floor of the chamber contained a white surface and was wiped with 70% ethanol before and after each use. Data were analysed by load cell recording (PanLab, Spain) and by video tracking (AnyMaze, Stoelting) with consistent results. The standard chamber context (Context A) included square arena (35 × 35 × 35 cm), blank wall and indirect red light (LED). Context was changed in experiments where specifically indicated to Context B (methacrylate walls, silver floor).

##### Habituation

Mice were habituated 24hrs to the conditioning chamber prior to fear conditioning (**Fig. 1f-g**). During habituation percent activity and freezing was measured and mice that displayed spontaneous freezing behaviour were excluded from further experimentation.

##### Entrainment

Mice were entrained using cued fear conditioning paradigm as previously described ^11^ with an optimised entrainment protocol (cue exposure length, shock amperage/length, conditioning context) to minimise effects of *tau* deficiency on LTM recall (at 24-48 hours) as reported with protocols of harsher entrainment ^24,48,58^. Entrainment sessions consisted of 180s habituation, followed by three auditory cue (CS) presentations (2kHz, 75dB) of 20s duration delivered at 180s, 260s and 340s and co-terminated with a 2s 0.6mA foot shock (US). Each tone-shock pairing consisted of a 60s intertrial interval. Following entrainment mice were returned to their home cage.

##### Recall testing

All recall test sessions were 420s in duration and consisted of 180s habituation followed by three tone presentations (2kHz, 75db) of 20s duration with no foot shock (-US). Memory was assessed for LTM at 24hrs and for ETM at 21-days and 42-days post-conditioning. We observe enhanced ETM fear response in wildtype mice due to reconsolidation of aversive memory associations by the single CS exposure during LTM ^59^, which has been reported previously for this conditioning paradigm ^5^.

##### Extinction trials

Extinction trials were performed as described previously ^60^. To assess active extinction by consecutive unpaired CS presentations, mice were habituated and entrained as for the standard CFC protocol above. 24 hours post-entrainment, mice were tested for LTM recall with unpaired CS (20s duration, 2kHz, 74dB) followed 24 hours later by presentation with 5 consecutive unpaired CS (120s duration, 2kHz, 74dB) within 10 minutes and freezing response was recorded and quantified.

##### Quantification of freezing behaviour

All behavioural experiments were analysed blind to experimental group. Behavioural data was recorded by load cell coupling (PanLab) and confirmed by video tracking (AnyMaze, Stoelting). For habituation and tone recall sessions, data were quantified using 5s freeze detection. Average freezing response was mapped onto chamber arena a custom script based on Rtrack ^61^ in RStudio (version 2023.06.0+421).

#### Touchscreen operant testing

Pairwise visual discrimination (VD) training was performed on a touchscreen operant chamber (Campden) as described previously ^16^ and based on developer’s protocols ^20^. Appetitive motivation was provided as liquid reward (Strawberry Iced Milk, Nippy’s). Priming of sessions at the start of each session provided an initial reward delivery (20µl). Mice used the same operant box for each session. The chamber interior was cleaned and wiped with 70% ethanol in between mice to equally distribute scents.

In brief, mice were placed on food restriction (2g regular chow/10 g bodyweight and day) and weight was monitored daily. Mice with > 10% loss of body weight were excluded from further experimentation. Food-restricted mice were habituated to the operant chamber on two consecutive days for 1 hour. Mice then underwent 4 stages of pre-training as described previously ^20^. For VD training, mice were presented with two stimuli (CS+ and CS-), for which reward association was randomised between subjects and pseudo-randomised location of stimuli. Correct (CS+) responses resulted in reward delivery (20µl) and tone (1s, 3 kHz), while incorrect choices (CS-) were followed by a correction trial (CT) with the same stimuli presentation after activated house light (5s) and an inter-trial interval (ITI). ITIs were defined at 20s. Mice were administered 1 hour of training daily at the same time of day with a maximum of 36 trials per session. For sufficient entrainment of VD acquisition, we defined a criterion of >80% correct trials in the last 2 consecutive acquisition sessions. Response efficiency was defined as correct choice count over the total session trials/time to complete the session trials, which takes into account accuracy and velocity of responses. Two months after the last acquisition session, mice were reintroduced into the operant chamber for a 1-day habituation session followed by retraining of operant behaviour for 4 consecutive sessions, prior to reacquisition of VD paradigm using the same CS+ and CS-pairings with randomisation as per initial acquisition to test for recall of learned CS+ associations. Recall efficiency was defined as average response efficiency for sessions 5-10.

#### Morris water maze

Spatial learning and memory was assessed in the Morris water maze as described previously ^16^ based on a standard protocol ^18^, with modifications to optimise for assessment of extended term memory. This paradigm is suitable for spatial acquisition in mice for ages up to 16 months to induce robust spatial memory ^15,45,62,63^. Mice in our study were of substantially lower age (6 months at ETM and reacquisition). The water maze tank was 145cm in diameter with white opaque high-density polyethylene walls (custom-built, Acrilix, Australia). The tank was filled with opacified room temperature water until 30cm from top edge and was surrounded equidistantly by off-white walls and partitions in an environment without high-contrast features on walls or ceiling and illuminated indirectly with low light. A hidden platform (12.6cm in diameter, custom-built) was located 1 cm below the water surface in quadrant opposite entry point for the initial spatial acquisition. Four off-white cue boards with salient visual cues (black geometric shapes; triangle, square circle, asterisk) were affixed at equidistant poles along the maze perimeter. Activity and movement were recorded with a ceiling-mounted digital camera (The Imaging Source) and analysed using automated tracking system (ANY-maze, Version 7.0, Stoelting Co, IL USA). Escape was defined as finding and ascending the hidden platform. Mice were habituated to the testing room and to the experimenter handling for 1 hr/day for 2-3 consecutive days prior to onset of spatial training. Entry into maze was semi-randomized at four positions along the perimeter of the entry quadrant. Mice were lowered into water maze facing the maze wall. For the initial *spatial acquisition*, the hidden platform position was kept constant for all mice. The initial acquisition phase (4 trials per day, 60sec per trial) was performed until mice reached spatial criterion, defined as <=15 sec on average for the entire cohort and <=20 second average of 4 trials, which resulted in acquisition phase of 1-6 days. Mice that did not reach criterion were excluded from recall trials due to lack of spatial acquisition. Spatial acquisition strategies were categorised as previously described ^64^. For non-spatial acquisition, the escape platform location was randomised between individual trials. For *spatial recall* at LTM and ETM two probe trials of 30s duration without escape platform were performed 24 hours and 4-months (120 days) after the last day of spatial acquisition, respectively. To evaluate reference memory for the spatial solution, a probe trial was administered 4-months (ETM) post-acquisition. To limit memory extinction due to lack of reinforcer (escape platform) during probe trials, the LTM probe trial was administered following 4-days of reacquisition after the ETM probe trial. Finally, mice underwent reversal learning and visual acuity testing (**Fig. 1j**). Spatial acquisition for LTM and ETM were towards hidden platforms in different quadrant annuli to avoid spatial bias in recall tests. LTM trials were administered 24 hours after the last reacquisition day to avoid the robust extinction effect on memory by void solution (i.e., probe) trials in the water maze ^18,65,66^. Recall trials were analysed (*1*) by tracking and quantifying crossings of the four quadrant annuli ^67^, (*2*) by cumulative time spent in the target quadrant ^18^, and (*3*) by occupancy mapping of location positions for each mouse per video frame onto the maze using a custom script based on Rtrack ^61^ in RStudio (version 2023.06.0+421). 24 hrs following ETM recall trial, mice were subjected to *spatial reacquisition* of the initial platform position until they reached criterion (3 1-min trials per session), followed by a reversal phase. In the *reversal* phase of testing, the ability of a mouse to learn a new location of the hidden platform after cue randomisation was assessed, which requires adaptation of allocentric and egocentric navigation modes in all individuals ^26,68^. For *visual cued trials*, a marker was attached to the hidden platform and escape latency recorded.

#### Open-field test

Open-field paradigm was done as previously described ^69^. Briefly, tests were performed in 40 × 40 × 40 cm square grey opaque Perspex boxes. Mice were placed in the arena facing the center and allowed 10 min for spontaneous exploration. Automated movement tracking was performed with ANY-maze (Stoelting). Time spent by mice in inner and outer zones of the open-field arena was quantified.

#### Elevated plus maze (EPM)

EPM tests were performed as previously described ^69^. Briefly, a custom-built EPM (dimensions: length of arms 35 cm; width of arms 5.5 cm; centre 5.5 × 5.5 cm^2^) with solid Perspex walls on closed arms (CA) and unwalled open arms (OA) was recorded above the maze centre and illuminated with bright indirect light. Lighting conditions were tested on an independent cohort to induce adequate anxiety-related response without complete suppression of explorative behaviour ^69^. Mice were acclimatized to the test room for 1 hour prior to testing. Mice were placed individually on the central platform of the EPM facing an open arm and their behaviour was recorded for 5 minutes. Videos were analysed using the AnyMaze software for the time spent exploring the CA vs. OA of the apparatus.

### Stereotactic injection to the hippocampus

All stereotactic intracranial infusion of viral vectors were performed following our previously published method ^44^. In brief, mice were anaesthetized using ketamine/xylazine (80/8 mg/kg of body weight, i.p.) and general anaesthesia was confirmed. The scalp was shaved and disinfected using alcoholic chlorhexidine. A midline incision was made, skull surface cleared using a retractor and mouse mounted onto the stereotaxic frame (Kopf Instruments, Tujunga, USA, catalogue nos. 940 and 926) with an animal temperature controller (TCAT-2, Physitemp, Clifton, USA) on a custom-built stage. The stereotaxic frame was centred on bregma as the reference position. Stereotaxic coordinates for bilateral hippocampal infusion (anterior-posterior (AP). -2.0; lateral (L), ±1.85; dorso-ventral (DV), −2.0] and AP, -3.6; ±2.5; DV, -3.2), were obtained from mouse brain atlas (Paxinos and Franklin, Fifth edition). Targeting the hippocampus is supported by its critical role for remote memory formation in spatial and associative learning ^26,27^, consistent with an indexing function associated with the hippocampal ensemble activated during encoding ^70^. Bur holes were made using a microdrill (Stoelting, #58610v) fitted with a 0.05-mm-diameter tungsten carbide drill head (Fine Science Tools, Meisinger #310104001). AAV-containing solution was infused using a 34-gauge needle (NF34BV-2, WPI) connected to a 10µL Nanofil Syringe (NANOFIL, WPI) attached to a precision nanolitre pump (UMP3, WPI) with controller (Micro4, WPI). The needle was lowered to target coordinates and remained for 2 minutes before infusion at 200nl/min. The injection needle was left in position for a further 5 minutes post volume delivery before retraction. After withdrawing of the needle, the incision was closed with staples and skin adhesive (Vetbond, 3M). Mice received analgesic (Buprenorphine (Temvet) 0.03mg/kg, s.c.) on the first recovery day and were allowed to recover for 1-week before subsequent experimentation.

### Doxycycline feed schedule for engram labelling

Doxycycline feeding schedule to regulate d2tTA-controlled expression from viral vectors (i.e., pAAV-*syn1*-d2tTA-eGFP/tau, pAAV-*RAM*-d2tTA-eGFP/YFP/tau) was performed based on previous protocols ^39^. For spatial memory engram labelling, mice were placed on doxycycline-containing diet (40mg/kg, Specialty Feeds) 5-7 days prior to stereotaxic injection of AAV containing a TRE::d2tTA expression cassette into the hippocampus and kept on DOX diet for 7 days post injection. DOX diet was removed 24 hrs prior to memory entrainment and replaced with regular chow (Gordons). Mice were either sacrificed 2 hours after the last stimulus presentation for histological assessment or placed on DOX diet again (200mg/kg, Specialty Feeds) and housed until memory recall and spatial reacquisition. For engram labelling of fear memory associations, all mice remained on DOX during the habituation phase. Following habituation DOX diet was removed and replaced by chow diet 24 hrs prior to entrainment and mice were kept on regular chow (-DOX). When entrainment was complete, mice were placed back on DOX-containing diet (200mg/kg, Specialty Feeds) until LTM and ETM testing.

### Telemetric Optogenetic stimulation of fear engram

Optogenetic targeting and stimulation of the fear engram was based on previous protocols ^5,71^. *Light cannula implantation* – Cannula implantation was performed following previously described methods ^72^. The telemetric optogenetic emitters (TeleR-2-P, TeleOpto, Japan) were equipped with bilateral LED cannulae (TeleLCD-B, TeleOpto, Japan; fiber diameter, 500 µm; fiber length, 1.8 mm; bilateral 3.8 mm distance, 473 nm) customised to target the mouse hippocampus above the DG. LED cannulae were implanted in a stereotaxic procedure under anaesthesia using ketamine/xylazine (80/8 mg/kg of body weight, i.p.) with the mouse mounted onto a stereotaxic frame (Kopf Instruments, Tujunga, USA, catalogue nos. 940 and 926) with temperature controller (TCAT-2, Physitemp, Clifton, USA). Prior to LED cannula implantation, hippocampus was dually infused with 50nl opsin-encoding AAV 50nl (pAAV-EF1a-double floxed-hChR2(H134R)-EYFP-WPRE-HGHpA (capsid PHP.B, 1.14 × 10^14^ viral genomes (vg)/ml) combined with 50nl activity-dependent doxycycline-regulated Cre recombinase-expressing virus (pAAV-*RAM*-d2tTA-Cre-WPRE-pA, capsid PHP.B, 8.14 × 10^12^ vg/ml) bilaterally. pAAV-EF1a-double floxed-hChR2(H134R)-EYFP-WPRE-HGHpA was a gift from Karl Deisseroth (Addgene plasmid # 20298 ; http://n2t.net/addgene:20298 ; RRID:Addgene_20298)^73^. After complete infusion and 5 minute incubation, the bone surface was scored and 4 anchors were installed to secure the cannula implant. The implant was lowered to the stereotaxic coordinates AP -2.0, L ±1.9, DV -1.8 and secured with acrylate. A cap of opaque dental cement (SpeedCEM, Ivoclar) was formed and cured. The scalp was closed, disinfected and the implant cap edges were covered with silicone cure (KwikSil, WPI). Mice were provided analgesic (TemVet, 0.03 mg/kg, s.c.) during recovery and experimentation was continued after full recovery. Mice were habituated to the emitter using a dummy 2-3 days prior to stimulation for 1 hour per day. *Stimulation protocol* – Entrainment, LTM and ETM with natural stimulus were all performed in the standard context (context A) as described for cued fear conditioning. Light stimulation to reactivate ChR2-expressing memory ensembles was performed in Context B (rounds arena) 24 hours after ETM recall testing. Freezing response was recorded in context B with and without light (473nm) stimulation (20Hz, 15ms, at 0.9-1.0 mW) based on comparable stimulation protocols of hippocampal engram ensembles ^5^. Light intensity at the cannula tip was confirmed pre- and post-experiments using a light power meter (LPM-100, TeleOpto, Japan).

### Telemetric electroencephalography

Mice were implanted with telemetry unilaterally into the hippocampus as previously ^74^. Briefly, after anaesthesia with ketamine/xylazine, a scalp incision along the midline was performed. The head was fixed in a stereotactic frame (Kopf instruments) and bregma was located. Bore holes were formed using a bone micro-drill (Fine Science Tools, F.S.T.) at positions previously described for the hippocampus (from bregma: -2.0 mm AP, ±2.0 mm ML, -2 mm DV). Electrodes were inserted at this position with reference electrode placed above the cerebellum (from bregma: -6.0 mm AP, 0 mm ML, 0 mm DV). Electrodes were fixed in place by polyacrylate followed by wound closure and rehydration. After complete recovery, local field potentials (LFP) were recorded at a sampling rate of 1000/s with a wireless receiver setup (DSI) with amplifier matrices using Ponemah software (DSI). Raw EEG signal was exported for further analysis. Correct placement of electrodes was confirmed by serial sections of paraffin embedded brain tissue with hematoxylin-eosin staining. Only recordings from mice with proper placement of electrodes were included in further analysis. Two days after the last EEG recordings were performed, animals were sacrificed by transcardial perfusion with cold phosphate-buffered saline (PBS) and brain samples were extracted for further processing for histological analysis.

Spectrogram analysis was performed using the spectrogram function in MATLAB (R2021a) in a custom script using 60 second data bins for each recording day in wake state. Cross-frequency coupling analysis was performed as previously ^16^ based on the method developed by Tort et al. ^31^ using a customised script in MATLAB (R2021a) and 90 second data bins in wake state.

### Molecular cloning

Sequences for eGFP, eYFP or tau (441 amino acids) as wildtype or T205A variant were amplified by polymerase chain reaction (PCR) using Q5 High-Fidelity master mix (M0492, New England Biolabs (NEB)) from previous plasmid constructs ^16^. PCR products were cloned into target constructs using HiFi Master mix (E2621, NEB). All constructs were verified by dideoxynucleotide sequencing (Macrogen), propagated in NEB Stable Competent *E.coli* (C3040, NEB) and isolated using endotoxin-free plasmid purification kit (Thermo). Addgene plasmids: Plasmids pAAV-*RAM*-d2TTA::TRE-MCS-WPRE-pA (Addgene# 63931) and pAAV-*RAM*-d2TTA::TRE-eGFP-WPRE-pA (Addgene# 84469) were a kind gift from Yingxi Lin. To generate doxycycline-regulated neuron-specific AAV expression constructs (pAAV-*syn1*-d2tTA::TRE-MCS-WPRE-pA), the RAM promoter in pAAV-*RAM*-d2TTA::TRE-MCS-WPRE-pA (Addgene# 63931) was replaced by a human *synapsin-1* promoter. Oligonucleotide primers for all cloning reactions are listed in Supplementary Table 3.

### Production of recombinant adeno-associated viral vectors

Packaging of AAV vectors was performed as previously ^44^. Briefly 293T cells were seeded in complete DMEM (Gibco, #11965175), supplemented with 10% FBS (Gibco, #10099), 1% (v/v) penicillin/streptomycin (Gibco, #10378016), and 2 mM GlutaMAX (L-alanyl-L-glutamine dipeptide supplement, Gibco, #35050061) at 70 to 80% confluence. Cell culture medium was replaced with Iscove’s modified Dulbecco’s medium (Gibco, #12240053) with 5% FBS (Gibco, #10099) three hours before transfection. Transfection included the viral genome-containing plasmid, pFΔ6 as helper plasmid, and AAV-PHP.B plasmid, encoding rep and cap sequences together with transfection reagent polyethylenimine-Max (PEI-Max; Polysciences, #24765-1) at a ratio of plasmid DNA:PEI of 3:1. Seventy-two hours following transfection, cells and supernatant were collected and clarified using 40% PEG 8000/2.5 M sodium chloride to a final concentration of 8% PEG 8000/0.5 M sodium chloride and incubated at 4°C for at least 2 hours. The cleared supernatant was spun at 2000*g* for 30 min at 4°C and combined cell pellet and precipitate containing viral particles were treated with sodium deoxycholate (Sigma-Aldrich, #D6750) and benzonase (∼500 U; Sigma-Aldrich, E8263) at 37°C for 40 min. Sodium chloride was added and incubated at 56°C for 40 min before cycles of freeze-thawing. The solution containing viral particles was spun for 30 min at 5000*g* and 4°C and purification of supernatants was performed by iodixanol (OptiPrep, Sigma-Aldrich, #D1556) gradient ultracentrifugation (475,900*g* for 2 hours at 18°C). Viral particles were concentrated using 100-kDa centrifugation filter units (Millipore, #ACS510024) at 5,000*g* and 4°C following buffer-exchange in PBS (pH 7.4) with the inclusion of non-ionic surfactant Pluronic F-68 (cat#. 24040032, Gibco) (final storage solution: PBS, 172 mM NaCl, 0.001% Pluronic F-68)). Titration of viral particles was performed by quantitative PCR (qPCR) using serial dilutions of purified AAV concentrates and Luna Universal qPCR mix (NEB#M3003). Oligonucleotide primers for AAV titration purpose were forward woodchuck hepatitis virus post-transcriptional regulatory element (WPRE) primer, 5′-GGCTGTTGGGCACTGACAAT-3’, and reverse WPRE primer, 5′-CCGAAGGGACGTAGCAGAAG-3’ (*<u>57</u>*). Determined AAV titres were AAV-*syn1*-d2tTA-eGFP (6.40 10^13^), AAV-*syn1*-d2tTA-tau (5.59 10^13^), AAV-*RAM*-d2tTA-eGFP (9.69 10^13^), AAV-*RAM*-d2tTA-eYFP (2.99 10^13^), AAV-*RAM*-d2tTA-tau (4.49 10^13^), AAV-*RAM*-d2tTA-tauT205A (2.48 10^13^), AAV-*RAM*-d2tTA-Cre (8.14 10^12^), pAAV-EF1a-double floxed-hChR2(H134R)-EYFP-WPRE-HGHpA, Addgene# 20298 (1.14 10^14^) (expressed as viral genomes per millilitre). AAV preparations were aliquoted and stored at −80°C.

### Tissue lysates

Lysates were prepared as previously described ^44^. Briefly, right murine cortical and/or hippocampal brain tissue was extracted upon transcardial perfusion with 1x PBS (pH 7.4) (Gibco, #10010023). Extracted tissue were immediately snap-frozen in liquid nitrogen for subsequent storage at −80°C. Tissue material was weighed before addition of ice-cold radioimmunoprecipitation assay (RIPA) buffer [20 mM tris (pH 8.0), 150 mM sodium chloride, 1 mM sodium EDTA, 1 mM activated sodium orthovanadate (Na_3_VO_4_), 5 mM NaF, 1 mM glycerophosphate, 2.5 mM sodium pyrophosphate (Na_2_H_2_P_2_O_7_), 1 mM phenylmethylsulfonyl fluoride, protease inhibitors (cOmplete, Roche, catalog no. 11697498001), 1% NP-40 substitute (Sigma-Aldrich, Merck, Munich, Germany, catalog no. 11754599001), 0.1% SDS, and 0.5% sodium deoxycholate] was added at a ratio of 10μl or 20μl of buffer per milligram hippocampal or cortical tissue weight respectively, and tissue was homogenized with a dounce homogenizer (10 strokes, 650 rpm; Heidolph, Schwabach, Germany) on ice. Resulting lysates were sonicated (2 × 2 s; 20% amplitude; Qsonica) before centrifugation at 16,000*g* for 10 min at 4**°**C. Protein concentrations were determined by colorimetric bicinchoninic acid (BCA) (Pierce, Thermo Fisher Scientific, cat# 23227).

### Immunoblots

To analyze protein expression, immunoblotting was performed as described previously ^44^. Briefly, tissue lysates (5µg total protein for AAV-*syn1*-d2tTA injected tissue; 30µg for AAV-*RAM*-d2tTA; 0.5µg for controls) were separated by SDS-PAGE (8 or 10%), transferred to nitrocellulose filter membranes (Immobilon-NC, Merck Millipore; catalog no. HATF00010 or Amersham Protran 0.45NC, GE10600002, Cytiva). Membranes were blocked in 3% milk powder in tris-buffered saline buffer with Tween 20 (TBS-T; 0.5%) at ambient temperature for 1 hr with gentle rocking. Primary antibodies were diluted in 3% bovine serum albumin (BSA, 0.05% sodium azide) in TBS-T overnight at 4°C with gentle rocking. Primary antibody was removed, membranes washed 3x 5minutes in TBS-T before incubation with Horseradish peroxidase (HRP)–coupled secondary antibodies for 1 hr at ambient temperature. HRP-coupled secondary antibodies used include goat–anti-rabbit (1:5000; Santa Cruz Biotechnology, #sc-2004) and goat–anti-mouse (1:5000; Santa Cruz Biotechnology, #sc-2005). Membranes were washed 3x 5 minutes before development of enhanced chemiluminescence reaction (BioRad). Chemiluminescence signals were imaged on a ChemiDoc MP (Bio-Rad) digital system. Primary antibodies used for immunoblots in this study were anti-tau (tau46) (1:5000; NEB 4019S), anti-human tau (tau13, 1:2000, Santa Cruz Biotechnology, #sc-21796), anti-tau D5D8N (1:1000; Cell Signaling Technology, #43894), anti-tau (1:4,000, A0024, Dako Agilent), anti–phospho–threonine-181 tau (1:1000; Abcam, #ab75679,), anti–phospho–serine-202 (1:1000; Abcam, #ab108387), anti– phospho–threonine-205 (1:2000; Abcam, #ab4841), anti–phospho–threonine-212 (1:1000; Abcam, #ab4842), anti–phospho–threonine-231 (1:1000; Abcam, #ab194815), anti–phospho–serine-235 (1:1000; Abcam, #ab306640), anti–phospho–serine-396 (1:2000; Abcam, #ab109390), anti–phospho– serine-404 (1:1000; Abcam, #ab92676), anti–phospho–serine-422 (1:1000; Abcam, #ab79415), anti-GAPDH (1:5000; Millipore, #ab2302), anti–green fluorescence protein (GFP; 1:1000; Abcam, #ab290), and anti-HA7 (1:5000; Sigma-Aldrich, #H3663).

Densitometric analysis of immunoblot images was done as previously described ^44^. Briefly, densitometric quantification of immunoblot results was performed using ImageJ 2.0.0-rc-49/1.51d. The rectangle analysis tool in ImageJ 2.0.0-rc-49/1.51d was used to select the length and width of the lane to be measured, beginning at the first lane, and using the identical frame across all lanes. Following this, the density of a band was defined by measurement of the total intensity peak area within the reduced dimension representation of the lane. A numerical value for each lane was generated, which was deducted from the background value of the immunoblot. Following this, all blots were normalized to the loading control (GAPDH) and tau expression. Subsequently, all values were analysed relative to the control condition. Each dataset is an average of at least three to four biological replicates per genotype and/or treatment group.

### Immunofluorescence

Histology and immunostaining procedures were performed as described previously ^75^. Mice were transcardially perfused with 1x ice-cold PBS and brains extracted and incubated in 4% (w/v) paraformaldehyde (PFA) overnight at 4°C. Brains were transferred to PBS and tissue was processed in an Excelsior tissue processor (Thermo Fisher) followed by paraffin embedding. Brains were sectioned at 5 μm. Sections were rehydrated and equilibrated in PBS pH7.4 before immunostaining procedure. For cryosections, mice were perfused with 1x PBS and brains extracted and incubated in 4% PFA overnight at 4°C. The following day, brains were transferred to ice-cold 1x PBS pH7.4 and placed to rotate for 1 hr at 4°C. Brains were then placed in consecutive steps of 10% for 1 h, 20% for 1h, and finally 30% sucrose in PBS at 4°C overnight until with gently rotation. Cryosections were prepared at 20μm thickness. Slices were washed 3x in PBS pH7.4. For egr-1 immunostaining, heat-induced antigen retrieval was performed in citrate buffer (100mM, pH6.0) at 70°C for 20 minutes before cooling slides to room temperature. Each slice was placed in PBS + 0.3% Triton X-100 (PBS-T) for 5 minutes, before washes with PBS-T with 3% normal goat or horse serum (consistent with the host species of the secondary antibody) for 1 hr before incubating with primary antibody at 4°C for 24 hr. Slices were washed 3x 10 min each in PBS-T, followed by 1 hr incubation with secondary antibody. For fluorescence staining, sections were incubated with Alexa Fluor-conjugated goat or donkey secondary antibodies (Thermo) and 4′,6-diamidine-2′-phenylindole dihydrochloride (DAPI) for 1 h at room temperature. Following three 10-min washes in PBS-T at room temperature, slices were mounted on superfrost plus microscope slides (Thermo) using water-based mounting medium (Fluoromount-G, Thermo or slowfade Diamond Antifade, Life Technologies). Antibodies for immunofluorescence stainings were anti-tau (mouse), anti-pT205 tau (Rabbit), anti-GFP (ab13970, Abcam, Chicken, 1:2,000), anti-cFos (9F6, Cell Signaling Technologies, 1:500), anti-egr-1 (9F6, Cell Signaling Technologies, 1:3,600), β3-tubulin (D71G9, Cell Signaling Technologies, 1:400). Immunofluorescence was imaged digitally using an AX70 epifluorescence microscope (Olympus, Tokyo, Japan) equipped with a DP80 colour camera (Olympus) operating through cellSens software (Olympus) or a slide scanner VS-200 (Olympus, Tokyo, Japan). Images were digitally processed using ImageJ (ImageJ NIH). For co-labelling experiments, percentage of labelled hippocampal cells were based on number of c-fos^+^ or Egr-1^+^ cells counted in Image J (v1.53q) and normalized to the number of nuclei (DAPI^+^) in the dentate gyrus subfield. Cells were counted from 4 coronal slices per mouse (n5-9 mice per group). All cell counting was conducted bli nd to experimental group.

### Statistical analysis

For statistical analysis, GraphPad Prizm (v7.0c) was used. Comparisons of two experimental groups were performed with unpaired, two-tailed Student’s *t* test for data of equal variance and Kolmogorov-Smirnov test for data of dissimilar variance. For comparisons of more than two experimental groups, analysis of variance (ANOVA) was performed with multiple comparisons with post-hoc testing. Data are expressed as means ± S.E.M. unless stated otherwise in the figure legend. A complete summary of statistical test parameters and results for data in main and supplementary figures is shown in Supplementary Table 3.

## Notes

### Competing Interest Statement

A.I. is a co-inventor on a patent application related to targeting tau in Alzheimer's and other neurodegenerative diseases (Australian patent number APA#2016900764).

