## Supplementary information and materials for "Remote memory engrams are controlled by encoding-specific tau phosphorylation"

**Author(s)**

Kristie Stefanoska[1,*], Emmanuel Prikas[1], Yijun Lin[1], Renée Kosonen[1], and Arne Ittner[1,*]

**Complete name(s) of institution(s)**

[1] Flinders Health and Medical Research Institute, College of Medicine and Public Health, Flinders University, Adelaide, Australia

### Supplementary Tables

**Table 1.**

**Oligonucleotide primers used for genotyping of mouse strains.**

| Strain | Forward Primer (5'-3') | Reverse Primer (5'-3') |
|---|---|---|
| *p38γ* | TGGGCTGCGAAGGTAGAGGTG | GTGTCACGTGCTCAGGGCCTG |
| *Thy1.2*-Cre | GCGGTCTGGCAGTAAAAACTATC | GTGAAACAGCATTGCTGTCACTT |
| tauT205A_4ARMS_I | CAGCCCCGGCTCTCCCGTAG | CGCGAGCGACTGCCAGGAGT |
| tauT205A_4ARMS_O | TGTATCAAAGTGACAGAGCAGGAGTGATGC | TGGTGCTTCAGGTTCTCAGTAGAGCCAA |
| *tau*$^{WT}$ | CTCAGCATCCCACCTGTAAC | CCAGTTGTGTATGTCCACCC |
| *tau*$^{KO}$ | AAGTTCATCTGCACCACCG | TGCTCAGGTAGTGGTTGTCG |
| *tauEx1* | GCAATCACCTTCCCTCCATA | ATTCAACCCCCTCGAATTTT |

**Table 2.**

**Oligonucleotide primers for HiFi PCR cloning reactions and quantitative PCR for recombinant AAV titering.**

| Construct/target | Forward Primer (5'-3') | Reverse Primer (5'-3') |
|---|---|---|
| pAAV-*syn1*-d2tTA-MCS-WPRE-pA | CCTGAAAGAAGATCCCCTGCAGACCCCC | CCAGCGTCCGTGTCACCC |
| pAAV-*syn1*-d2tTA-eGFP-WPRE-pA | TCCCCTGCAGGAACCCACTGAGG | GATTCTTTCAGGCCAGCG |
| pAAV-*syn1*-d2tTA-tau-WPRE-pA | TCCCCTGCAGGCACCCACTGAGG | GATTCTTTCAGGCCAGCG |
| pAAV-*RAM*-d2tTA-eGFP-WPRE-pA | TGCTAAGAGCGAACCAACAGCGG | TCAGAGGTTTCAGAGCCC |
| pAAV-*RAM*-d2tTA-tau(WT/T205A)-WPRE-pA | TGCTAAGAGCGCACCAACAGCGG | TCAGAGGTTTCAGAGCCC |
| pAAV-*RAM*-d2tTA-Cre-WPRE-pA | TCGCCACCGGTGTTTAAACGATGCCCAAGAAGAAGAGG | GGTGGATCGCTGGCGCGCCGCTAATCGCCATCTTCCAGCAG |
| WPRE qPCR | CATTGGAGACGAACCCAGCCTGG | CCTGCTTCTTCAGCTGTG |

**Table 3. Statistical information**

Details of statistical analyses used for each experimental condition in main Figures 1 to 4. P values and n numbers are provided in individual figure legends. Statistical analysis was done with Prism 6 (GraphPad). Abbreviations: GT, genotype; AAV, AAV-delivered transgene

| Main figure | Test |
|---|---|
| 1c | Unpaired *t*-test, two-tailed: *t*=1.160; *df*=49; α=0.05; n(WT/KO)=27/24 |
| 1d | Unpaired *t*-test, two-tailed: *t*=3.594; *df*=49; α=0.05; n(WT/KO)=27/24 |
| 1g | Two-way ANOVA: $F_{GT}$(1, 180)=0.0027; $F_{Sessions}$(8, 180)=11.73; α=0.05; Šídák post-hoc; n($tau^{+/+}$/$tau^{-/-}$)=9/13 |
| 1h | Unpaired *t*-test, two-tailed: *t*=3.397; *df*=118; α=0.05; n($tau^{+/+}$/$tau^{-/-}$)=9/13 |
| 1j | Two-way ANOVA: $F_{GT}$(1, 133)=1.535; $F_{Day}$(1, 133)=20.92; α=0.05; Tukey post-hoc; n($tau^{+/+}$/$tau^{-/-}$)=15/20 |
| 1l | Unpaired *t*-test, two-tailed: *t*=0.1532; *df*=33; α=0.05; n($tau^{+/+}$/$tau^{-/-}$)=15/20 |
| 1m | Unpaired *t*-test, two-tailed: *t*=4.218; *df*=33; α=0.05; n($tau^{+/+}$/$tau^{-/-}$)=15/20 |
| 1n | Two-way ANOVA: $F_{GT}$(1, 201)=17.89; $F_{Day}$(6, 201)=9.011; α=0.05; Šídák post-hoc; n($tau^{+/+}$/$tau^{-/-}$)=15/20 |
| 2c’ | One-way ANOVA: F(3, 36)=21.59; α=0.05; Šídák post-hoc; n=8-12 |
| 2e | Two-way ANOVA: $F_{GT}$(1, 308)=0.0753; $F_{Day}$(1, 308)=0.0753; α=0.05; Šídák post-hoc; n(syn-d2tTA-eGFP/syn-d2tTA-tau)=22/23 |
| 2g | Unpaired *t*-test, two-tailed: *t*=0.6338; *df*=43; α=0.05; n(syn-d2tTA-eGFP/syn-d2tTA-tau)=22/23 |
| 2h | Unpaired *t*-test, two-tailed: *t*=3.157; *df*=43; α=0.05; n(syn-d2tTA-eGFP/syn-d2tTA-tau)=22/23 |
| 2i | Two-way ANOVA: $F_{AAV}$(1, 161)=9.157; $F_{Day}$(6, 161)=6.555; α=0.05; Šídák post-hoc; n(syn-d2tTA-eGFP/syn-d2tTA-tau)=12/13 |
| 2k | Unpaired *t*-test, two-tailed: *t*=1.005; *df*=41; α=0.05; n(syn-d2tTA-eGFP/syn-d2tTA-tau)=21/22 |
| 2l | Unpaired *t*-test, two-tailed: *t*=2.866; *df*=41; α=0.05; n(syn-d2tTA-eGFP/syn-d2tTA-tau)=21/22 |
| 2p | Unpaired *t*-test, two-tailed: *t*=3.684; *df*=30; α=0.05; n($tau^{+/+}$/$tau^{-/-}$)=6/7 |
| 2q | Kolmogorov-Smirnov test: *D*=0.8975; α=0.05; n(syn-d2tTA-eGFP/syn-d2tTA-tau)=6/9 |
| 3a’ | One-way ANOVA: $F_{pT205}$(3, 26)=15.00; α=0.05; Tukey post-hoc; n=8-12 |
| 3f | Two-way ANOVA: $F_{GT}$(1, 180)=3.331; $F_{Day}$(5, 180)=51.98; α=0.05; Šídák post-hoc; n(*tau*$T205^{T/T}$/*tau*$T205^{A/A}$)=18/20 |
| 3h | Unpaired *t*-test, two-tailed: *t*=0.1788; *df*=36; α=0.05; n(*tau*$T205^{T/T}$/*tau*$T205^{A/A}$)=18/20 |
| 3i | Unpaired *t*-test, two-tailed: *t*=3.291; *df*=36; α=0.05; n(*tau*$T205^{T/T}$/*tau*$T205^{A/A}$)=18/20 |
| 3j | Two-way ANOVA: $F_{GT}$(1, 219)=26.07; $F_{Day}$(6, 219)=15.70; α=0.05; Šídák post-hoc; n(*tau*$T205^{T/T}$/*tau*$T205^{A/A}$)=18/20 |
| 3l | Unpaired *t*-test, two-tailed: *t*=0.4284; *df*=28; α=0.05; n(*tau*$T205^{T/T}$/*tau*$T205^{A/A}$)=15/16 |
| 3m | Unpaired *t*-test, two-tailed: *t*=4.685; *df*=31; α=0.05; n(*tau*$T205^{T/T}$/*tau*$T205^{A/A}$)=15/16 |
| 3q | Unpaired *t*-test, two-tailed: *t*=3.373; *df*=14; α=0.05; n(*tau*$T205^{T/T}$/*tau*$T205^{A/A}$)=5/5 |
| 4d’ | ANOVA: *W*=19.14; *Dfn*=2.000; Dfd=29.58 Welch post-hoc; n(RAM-d2tTA-eGFP/RAM-d2tTA-$tau^{WT}$/RAM-d2tTA-$tau^{T205A}$)=6/6/6 |
| 4f | Two-way ANOVA: $F_{GT}$(1, 162)=3.464; $F_{Day}$(4, 162)=36.4; α=0.05; Šídák post-hoc; n(RAM-d2tTA-$tau^{WT}$/RAM-d2tTA-$tau^{T205A}$)=14/15 |
| 4h | Unpaired *t*-test, two-tailed: *t*=0.6154; *df*=28; α=0.05; n(RAM-d2tTA-$tau^{WT}$/RAM-d2tTA-$tau^{T205A}$)=14/15 |
| 4i | Unpaired *t*-test, two-tailed: *t*=2.789; *df*=28; α=0.05; n(RAM-d2tTA-$tau^{WT}$/RAM-d2tTA-$tau^{T205A}$)=14/15 |
| 4j | Two-way ANOVA: $F_{GT}$(1, 231)=17.92; $F_{Day}$(6, 231)=13.52; α=0.05; Šídák post-hoc; n(RAM-d2tTA-$tau^{WT}$/RAM-d2tTA-$tau^{T205A}$)=14/15 |
| 4l | Unpaired *t*-test, two-tailed: *t*=1.669; *df*=32; α=0.05; n(RAM-d2tTA-$tau^{WT}$/RAM-d2tTA-$tau^{T205A}$)=16/18 |

| | |
|---|---|
| 4m | Unpaired *t*-test, two-tailed: *t*=3.928; *df*=32; α=0.05; n(RAM-d2tTA-tau$^{WT}$/RAM-d2tTA-tau$^{T205A}$)=16/18 |
| 4r | Unpaired *t*-test, two-tailed: *t*=0.070; *df*=15; α=0.05; n(WT/KO)=9/8 |
| 4s | Unpaired *t*-test, two-tailed: *t*=3.346; *df*=15; α=0.05; n(WT/KO)=9/8 |
| 4t | Two-way ANOVA: $F_{GT}$(1, 64)=0.742, $F_{Stimulation}$(3, 64)=19.87; α=0.05; Tukey's post-hoc; n(*tau*$^{+/+}$/*tau*$^{-/-}$)=9/8 |
| **Supplementary** | **figures** |
| S1a | Unpaired *t*-test, two-tailed: *t*=1.173; *df*=49; α=0.05, n(*tau*$^{+/+}$/*tau*$^{-/-}$)=27/24 |
| S1b | Unpaired *t*-test, two-tailed: *t*=0.8640; *df*=49; α=0.05, n(*tau*$^{+/+}$/*tau*$^{-/-}$)=27/24 |
| S1c | Unpaired *t*-test, two-tailed: *t*=1.923; *df*=49; α=0.05, n(*tau*$^{+/+}$/*tau*$^{-/-}$)=20/18 |
| S1d | Two-way ANOVA: $F_{Genotype}$(1, 368)=0.2302; $F_{Extinction}$(10, 368)=12.76; α=0.05; Šídák post-hoc test; n(*tau*$^{+/+}$/*tau*$^{-/-}$)=20/15 |
| S1e | Two-way ANOVA: $F_{Genotype}$(1, 77)<0.0001; $F_{Zone}$(1, 77)=133.1; α=0.05; Šídák post-hoc test; n(*tau*$^{+/+}$/*tau*$^{-/-}$)=20/15 |
| S1f | Two-way ANOVA: $F_{Genotype}$(1, 41)<0.0001; $F_{Arm}$(1, 41)=182.6; α=0.05; Šídák post-hoc test; n(*tau*$^{+/+}$/*tau*$^{-/-}$)=12/10 |
| S2b | Two-way ANOVA: $F_{GT}$(1, 160)=15.89; $F_{Sessions}$(9, 160)=6.088; α=0.05; Šídák post-hoc; n(*tau*$^{+/+}$/*tau*$^{-/-}$)=9/13 |
| S3b | Two-way ANOVA: $F_{GT}$(1, 175)=0.1069; $F_{Day}$(6, 175)=18.61;; α=0.05; Šídák post-hoc; n(*tau*$^{+/+}$/*tau*$^{-/-}$)=19/19 |
| S3c | Two-way ANOVA: $F_{GT}$(1, 144)=0.7177; $F_{Annulus}$(3, 144)=8.823; α=0.05; Šídák post-hoc; n(*tau*$^{+/+}$/*tau*$^{-/-}$)=19/19 |
| S3d | Two-way ANOVA: $F_{GT}$(1, 175)=0.1168; $F_{Day}$(6, 175)=10.44; α=0.05; Šídák post-hoc; n(*tau*$^{+/+}$/*tau*$^{-/-}$)=18/19 |
| S3e | Two-way ANOVA: $F_{GT}$(1, 140)=4.497; $F_{Annulus}$(3, 140)=19.32; α=0.05; Šídák post-hoc; n(*tau*$^{+/+}$/*tau*$^{-/-}$)=18/19 |
| S3g | Two-way ANOVA: $F_{GT}$(1, 128)=4.188; $F_{Annulus}$(3, 128)=1.539; α=0.05; Šídák post-hoc; n(*tau*$^{+/+}$/*tau*$^{-/-}$)=18/19 |
| S4b' | as 2c' |
| S5b | Two-way ANOVA: $F_{AAV}$(1, 152)=0.037; $F_{Annulus}$(3, 152)=22.49; α=0.05; Šídák post-hoc; n(syn-d2tTA-eGFP/syn-d2tTA-tau)=22/23 |
| S5c | Two-way ANOVA: $F_{AAV}$(1, 152)=9.840; $F_{Annulus}$(3, 152)=2.899; α=0.05; Šídák post-hoc; n(syn-d2tTA-eGFP/syn-d2tTA-tau)=22/23 |
| S6a | Unpaired *t*-test, two-tailed: *t*=0.826; *df*=41; α=0.05; n(syn-d2tTA-eGFP/syn-d2tTA-tau)=21/22 |
| S6b | Unpaired *t*-test, two-tailed: *t*<0.0001; *df*=41; α=0.05; n(syn-d2tTA-eGFP/syn-d2tTA-tau)=21/22 |
| S6c | Unpaired *t*-test, two-tailed: *t*=2.565; *df*=17; α=0.05; n(syn-d2tTA-eGFP/syn-d2tTA-tau)=10/9 |
| S8a | Unpaired *t*-test, two-tailed: *t*=0.032; *df*=30; α=0.05; n(*tau*$^{+/+}$/*tau*$^{-/-}$)=6/7 |
| S8b | Unpaired *t*-test, two-tailed: *t*=3.245; *df*=30; α=0.05; n(*tau*$^{+/+}$/*tau*$^{-/-}$)=6/7 |
| S8c | Unpaired *t*-test, two-tailed: *t*=2.025; *df*=30; α=0.05; n(*tau*$^{+/+}$/*tau*$^{-/-}$)=6/7 |
| S9a' | One-way ANOVA: $F_{tau}$(3, 36)=92.04; α=0.05; Šídák post-hoc; n=8-12 |
| S9a''' | One-way ANOVA: $F_{pT181}$(3, 36)=8.303; $F_{pS202}$(3, 26)=14.30; $F_{pT205}$(3, 26)=15.00; $F_{pT212}$(3, 12)=0.6037; $F_{pT217}$(3, 12)=8.303; $F_{pT231}$(3, 12)=0.431; $F_{pS235}$(3, 12)=0.337; $F_{pS396}$(3, 16)=0.475; $F_{pS404}$(3, 12)=3.014; α=0.05; Šídák post-hoc; n=3-12 |
| S10b | Two-way ANOVA: $F_{GT}$(1, 136)=3.834; $F_{Annulus}$(3, 136)=6.238; α=0.05; Šídák post-hoc; n(*tau*T205$^{T/T}$/*tau*T205$^{A/A}$)=18/20 |
| S11b | Two-way ANOVA: $F_{GT}$(1, 144)=3.834; $F_{Day}$(5, 144)=20.33; α=0.05; Šídák post-hoc; n(*p38γ*$^{+/+}$/ *p38γ*$^{-/-}$)=14/12 |
| S11c | Unpaired *t*-test, two-tailed: *t*=2.665; *df*=24; α=0.05; n(*p38γ*$^{+/+}$/ *p38γ*$^{-/-}$)=14/12 |
| S11d | Unpaired *t*-test, two-tailed: *t*=2.294; *df*=24; α=0.05; n(*p38γ*$^{+/+}$/ *p38γ*$^{-/-}$)=14/12 |
| S12a | Unpaired *t*-test, two-tailed: *t*=1.019; *df*=29; α=0.05; n(*tau*T205$^{T/T}$/*tau*T205$^{A/A}$)=15/16 |
| S12b | Unpaired *t*-test, two-tailed: *t*=1.019; *df*=29; α=0.05; n(*tau*T205$^{T/T}$/*tau*T205$^{A/A}$)=15/16 |
| S12c | Two-way ANOVA: $F_{GT}$(1, 198)=3.106; $F_{Extinction}$(10, 198)=7.369; α=0.05; Šídák post-hoc; n(*tau*T205$^{T/T}$/*tau*T205$^{A/A}$)=12/8 |

| | |
|---|---|
| S12d | Two-way ANOVA: $F_{GT}$(1, 36)=0.01; $F_{OpenA\text{-}ClosedA}$(3, 36)=229.9; α=0.05; Šídák post-hoc; n(*tau*T205$^{T/T}$/*tau*T205$^{A/A}$)=12/8 |
| S13a | Unpaired *t*-test, two-tailed: *t*=0.3916; *df*=14; α=0.05; n(*tau*T205$^{T/T}$/*tau*T205$^{A/A}$)=5/5 |
| S13b | Unpaired *t*-test, two-tailed: *t*=2.286; *df*=14; α=0.05; n(*tau*T205$^{T/T}$/*tau*T205$^{A/A}$)=5/5 |
| S13c | Unpaired *t*-test, two-tailed: *t*=1.582; *df*=14; α=0.05; n(*tau*T205$^{T/T}$/*tau*T205$^{A/A}$)=5/5 |
| S14a | ANOVA: $F_{tau}$(2, 9)=0.6145; α=0.05; Šídák post-hoc; n(RAM-d2tTA-eGFP/RAM-d2tTA-tau$^{WT}$/RAM-d2tTA-tau$^{T205A}$)=3/6/3 |
| S16b | Two-way ANOVA: $F_{GT}$(1, 144)=0.1710; $F_{Annulus}$(3, 144)=80.66; α=0.05; Šídák post-hoc; $t_{T205,\ A2}$=1.964; n(RAM-d2tTA-tau$^{WT}$/RAM-d2tTA-tau$^{T205A}$)=14/15 |
| S16c | Two-way ANOVA: $F_{GT}$(1, 144)=0.1437; $F_{Annulus}$(3, 144)=4.191; α=0.05; Šídák post-hoc; $t_{T205,\ A1}$=3.393; n(RAM-d2tTA-tau$^{WT}$/RAM-d2tTA-tau$^{T205A}$)=14/15 |
| S17b | Unpaired *t*-test, two-tailed: *t*=1.794; *df*=32; α=0.05; n(RAM-d2tTA-tau$^{WT}$/RAM-d2tTA-tau$^{T205A}$)=16/18 |
| S17c | Unpaired *t*-test, two-tailed: *t*=1.917; *df*=32; α=0.05; n(RAM-d2tTA-tau$^{WT}$/RAM-d2tTA-tau$^{T205A}$)=16/18 |
| S18a | Unpaired *t*-test, two-tailed: *t*=0.4308; *df*=11; α=0.05; n(*tau*$^{+/+}$/*tau*$^{-/-}$)=7/6 |

**Supplementary figures and figure legends**

**Supplementary Figure 1**

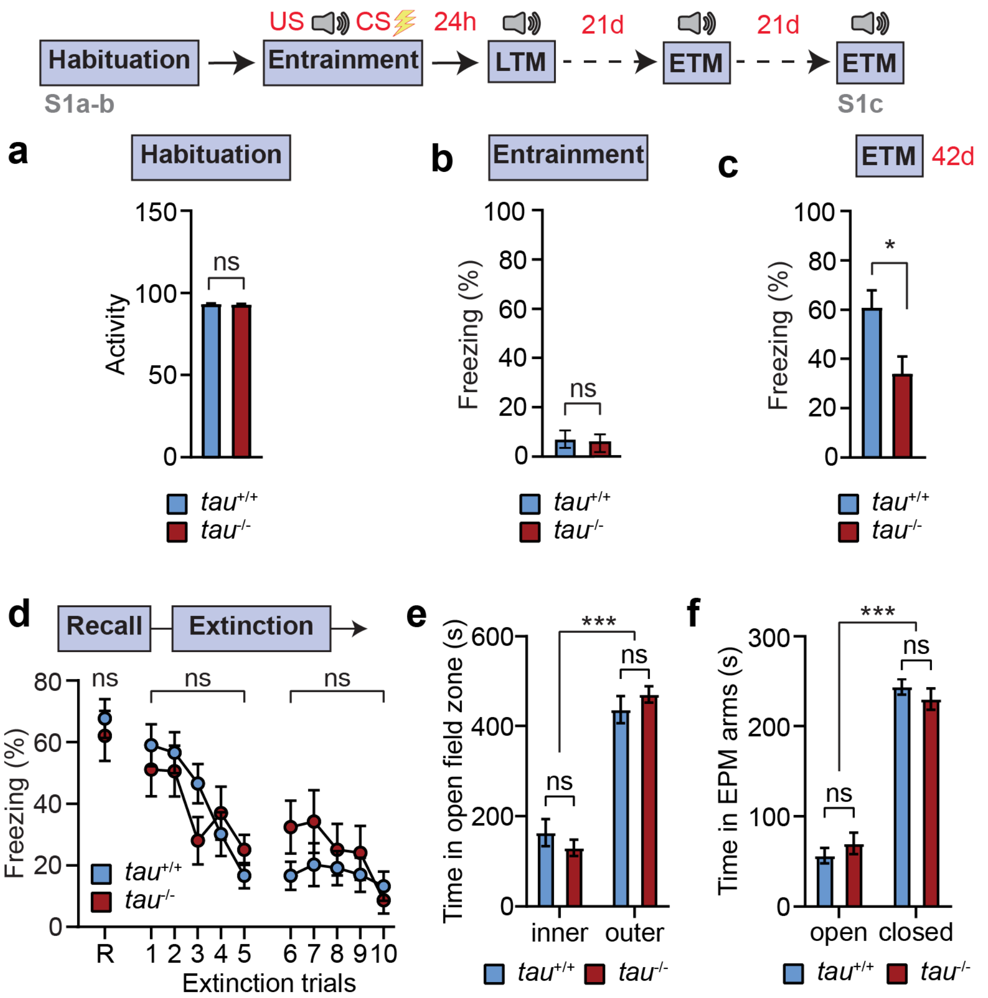

***Tau$^{-/-}$* mice show no enhanced trait nor state anxiety compared with *tau$^{+/+}$* mice.**

(**a**) Activity recorded during habituation phase of CFC schedule in *tau*$^{+/+}$ and *tau*$^{-/-}$ mice. No difference was observed indicating comparable trait anxiety and general activity levels in both genotypes. (n=17-23) ns, not significant (Student's t-test).

(**b**) Stimulus-induced freezing during entrainment phase in CFC schedule in *tau*$^{+/+}$ and *tau*$^{-/-}$ mice. No difference was observed indicating comparable state anxiety. (n=17-23) ns, not significant (Student's t-test).

(**c**) Extended-term memory (ETM) recall at 42 days post-conditioning in *tau*$^{+/+}$ and *tau*$^{-/-}$ mice (n=18-20). Note that these mice received a recall trial at 21days post-conditioning, which represents an extinction trial. At d42 ETM, *tau*$^{-/-}$ show significantly lower recall than *tau*$^{+/+}$.

(**d**) Extinction of fear memory in *tau*$^{+/+}$ and *tau*$^{-/-}$ mice entrained by cued fear conditioning (n=18-20). After entrainment and LTM recall (R), mice were presented with an unpaired CS (2 min) in 5 consecutive trials on each of 2 consecutive days (extinction trials 1-10). Freezing response was recorded for initial LTM recall (R) of fear memory and each extinction trial. Note that *tau*$^{-/-}$ mice show comparable freezing response during extinction trials as *tau*$^{+/+}$ mice, precluding a role of tau in active memory extinction.

(**e**) Spontaneous levels of inner zone and outer zone activity in the open field in *tau*$^{+/+}$ and *tau*$^{-/-}$ mice. (n=18-22) ns, not significant (Student's t-test). No difference was observed indicating comparable trait anxiety.

(**f**) Spontaneous occupancy of open and closed arms of the elevated plus maze (EPM) for *tau*$^{+/+}$ and *tau*$^{-/-}$ mice. (n=10-12)

Statistical comparisons are performed using ANOVA (d-f) or Student's t-test (a-c); ***$p<0.001$, *$p<0.05$; ns, not significant. Data are presented as mean ± S.E.M.

**Supplementary Figure 2**

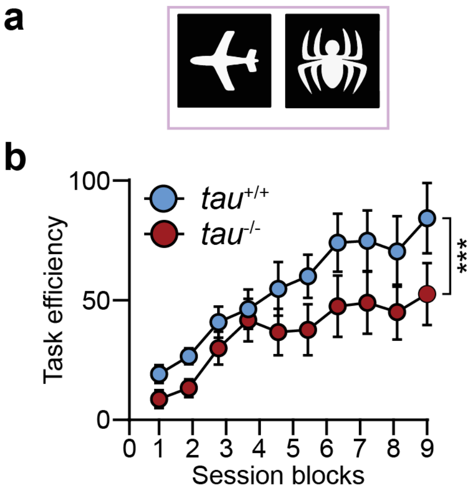

**Delayed reacquisition (after 2 months of initial training) in pairwise discrimination learning in *tau*$^{-/-}$ as compared with *tau*$^{+/+}$ mice.**

(**a**) Visual stimuli presented in the pairwise visual discrimination (VD) task. Stimulus position and appetitive CS (strawberry milkshake) pairing are randomised between mice and trials to avoid position and visual cue bias.

(**b**) Reacquisition of VD learning in *tau*$^{+/+}$ and *tau*$^{-/-}$ mice (n=8) 2 months after completion of initial VD acquisition sessions (see Fig. 1h). Note that *tau*$^{-/-}$ mice show persistent delay in task reacquisition across the 9 session blocks.

Statistical comparisons are performed using ANOVA; ***$p<0.001$. Data are presented as mean ± S.E.M.

**Supplementary Figure 3**

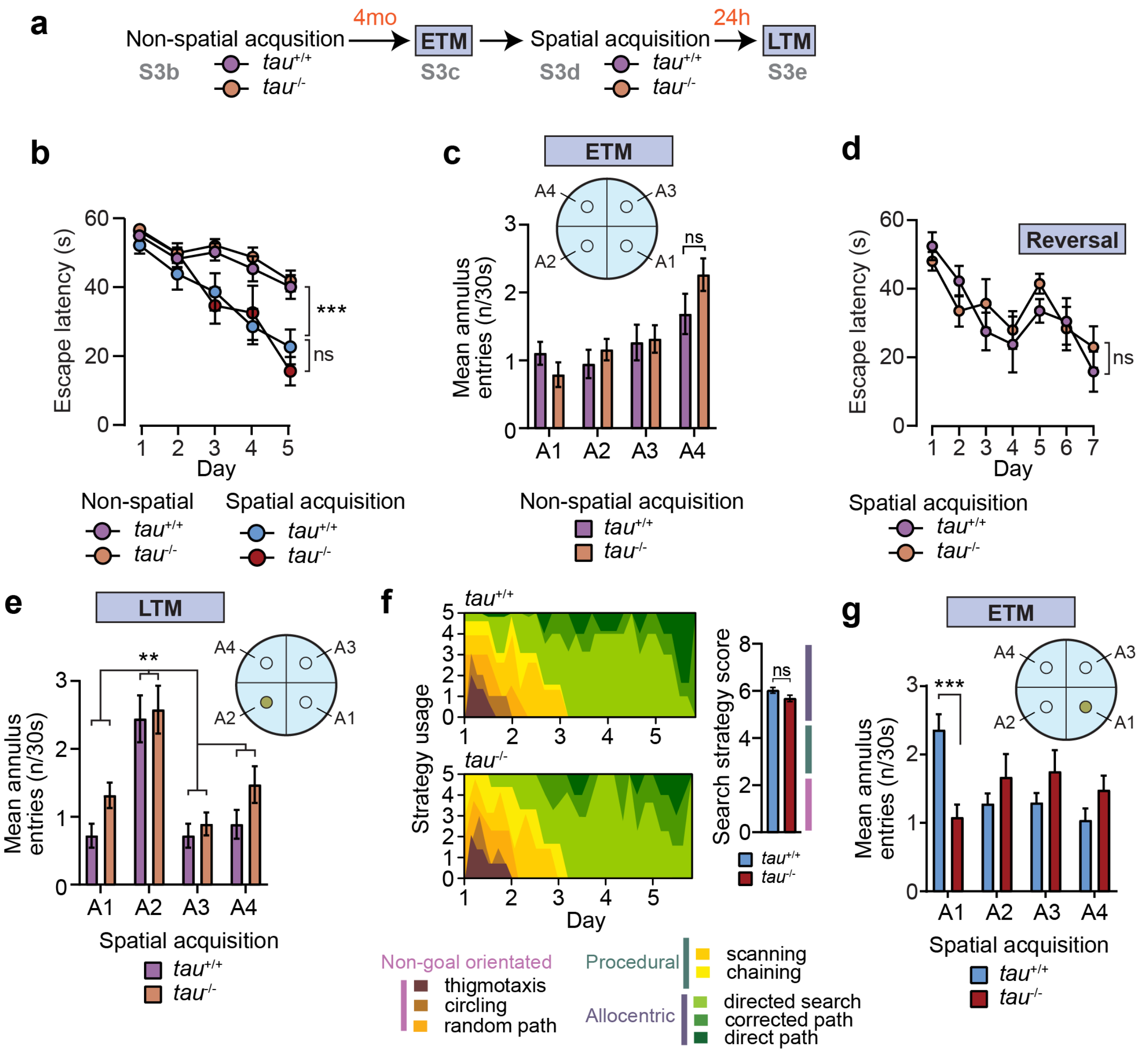

**Non-spatial acquisition in the Morris water maze (MWM) is unaltered in the absence of *tau***

(**a-e**) Comparison of spatial and non-spatial acquisition in the MWM in $tau^{+/+}$ and $tau^{-/-}$ mice (2-month-old). (n=18-19)

(**a**) Experimental schedule for $tau^{+/+}$ and $tau^{-/-}$ mice undergoing non-spatial acquisition with extended-term recall at 4 months post-non-spatial acquisition, subsequently followed by spatial acquisition and long-term recall on day 1 post-spatial acquisition.

(**b**) Acquisition curve in mice of indicated genotypes with random escape platform position at each trial in the absence of visual cues (= non-spatial) and with specific escape platform location in presence of visual cues (= spatial).

(**c**) ETM recall of *tau*$^{+/+}$ and *tau*$^{-/-}$ mice 4 months after non-spatial acquisition in MWM. Note that prior non-spatial acquisition induces a bias to cross position A4 near the maze entry point during each trial reflective of episodic memory of the acquisition trials as well as of the lack of escape solution.

(**d**) Spatial acquisition curve of *tau*$^{+/+}$ and *tau*$^{-/-}$ mice 4 months after non-spatial acquisition in MWM. Reversal with complete randomisation of cues and hidden platform position was done on day 5.

(**e**) LTM recall of *tau*$^{+/+}$ and *tau*$^{-/-}$ mice 24 hours after spatial acquisition in MWM with hidden platform in position A2.

(**f**) Spatial search strategy analysis in *tau*$^{+/+}$ and *tau*$^{-/-}$ mice in spatial acquisition in Figures 1k and Fig S3a. Search strategies were comparably employed by both *tau*$^{+/+}$ and *tau*$^{-/-}$ and gravitated towards allocentric strategies towards day 5 of the acquisition phase for both genotypes. Overall search strategy score was comparable between spatial acquisition in *tau*$^{+/+}$ and *tau*$^{-/-}$ mice.

(**g**) Annulus analysis of ETM recall in *tau*$^{+/+}$ and *tau*$^{-/-}$ with spatial acquisition 4 months prior. *Tau*$^{-/-}$ mice show significantly fewer entries into the target annulus A1 and more entries into non-target annuli A2-4 on average during ETM recall trials as compared with *tau*$^{+/+}$ mice. (n=15-20)

Statistical comparisons are performed using ANOVA (b-e, g) and Mann-Whitney rank test (f); ***$p<0.001$, **$p<0.01$; ns, not significant. Data are presented as mean ± S.E.M.

**Supplementary Figure 4**

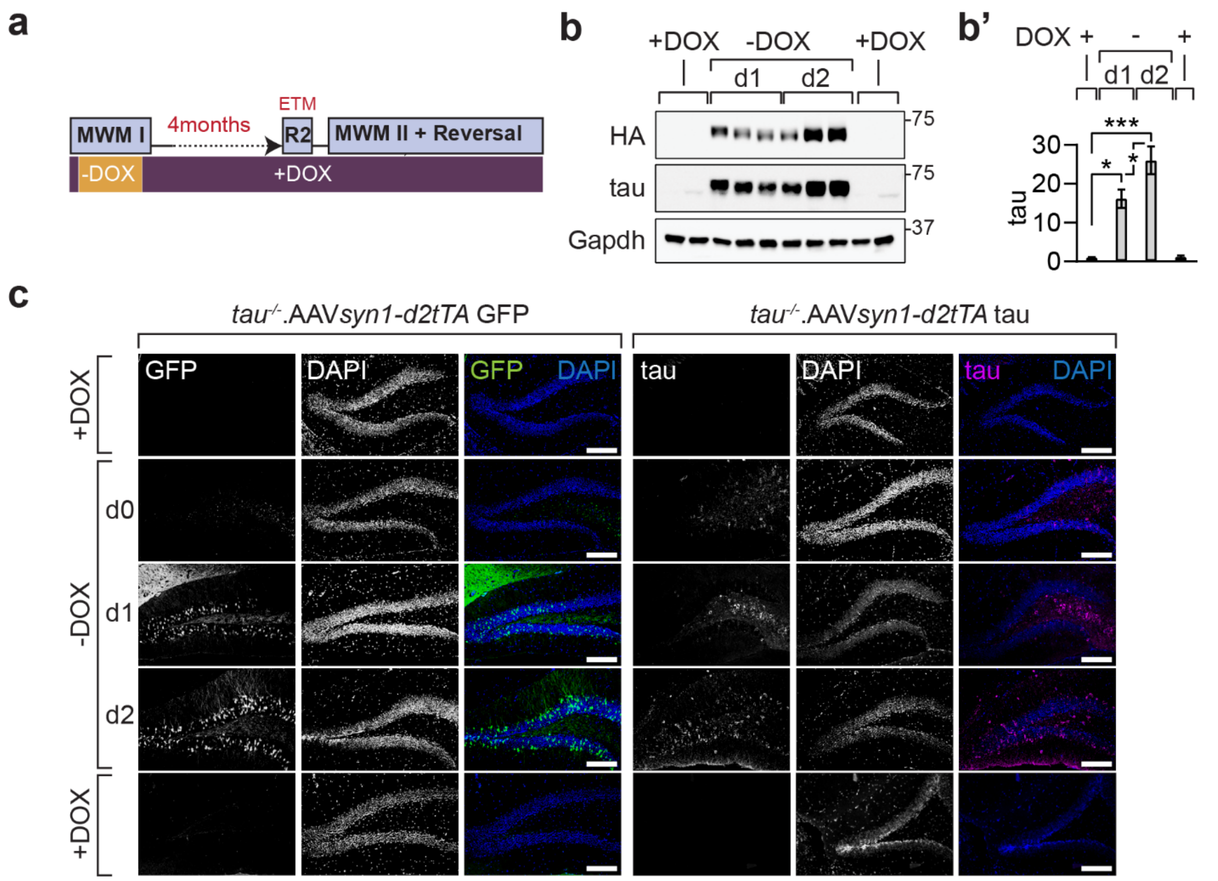

**Temporally controlled neuronal tau expression during spatial acquisition in the Morris water maze**

(**a**) Schematic of controlled expression from AAV *syn1*-d2tTA-tau or -eGFP vectors delivered to the hippocampus of *tau*$^{-/-}$ mice prior to acquisition in the Morris water maze (MWM I) with suppression of expression prior to and after spatial acquisition (+DOX, purple shade). Expression is induced by removal of doxycycline from diet (-DOX, orange shade) from day 0 to 3 of the spatial acquisition. Mice undergo ETM recall after 4 months of latency phase, followed by reacquisition (MWM II) and reversal learning.

(**b**) Representative expression analysis by immunoblot for tau from hippocampal lysates from *tau*$^{-/-}$. AAV *syn1*-d2tTA-tau mice with indicated DOX regime at indicated days of spatial acquisition in the MWM. Transgenic tau is fused to an N-terminal haemagglutinin (HA) tag. Gapdh, loading control for immunoblot. (**b'**) Quantification of immunoblots for controlled tau expression. (n=6) Tau levels are represented relative to low level signal with -DOX and relative to Gapdh to show the ~15-fold degree of induction.

(**c**) Immunofluorescence analysis of eGFP or tau expression in the hippocampal formation (dentate gyrus) of *tau*$^{-/-}$. AAV *syn1*-d2tTA-tau mice at indicated time points after DOX removal. D0, corresponds to 12 hours post-removal of DOX block. Signals for eGFP or tau are virtually absent with DOX administration in diet confirming dynamic regulation of d2tTA-dependent viral transgene expression. Scale bar, 200 µm.

Statistical comparisons are performed using ANOVA; ***$p<0.001$, *$p<0.05$; ns, not significant. Data are presented as mean ± S.E.M.

**Supplementary Figure 5**

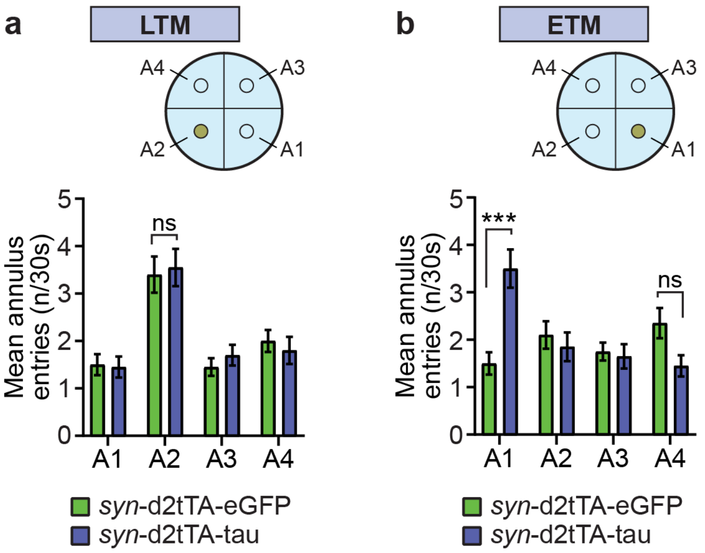

**Additional parameters of spatial learning and memory in mice with controlled neuronal tau expression**

(**a-b**) Annulus analysis of (**a**) LTM and (**b**) ETM recall in *tau*$^{+/+}$ and *tau*$^{-/-}$ with spatial acquisition 4 months prior. *Tau*$^{-/-}$ mice injected with AAV *syn1*-d2tTA-tau (with induced expression during spatial acquisition; *see Fig. 2a-d and fig S4*) show significantly more entries into the target annulus A1 and more entries into non-target annuli A2-4 on average during ETM recall trials as compared with AAV *syn1*-d2tTA-eGFP-injected controls. (n=22-23) *** $p < 0.001$; ns, not significant (ANOVA)

Statistical comparisons are performed using ANOVA; ***$p<0.001$; ns, not significant. Data are presented as mean ± S.E.M.

**Supplementary Figure 6**

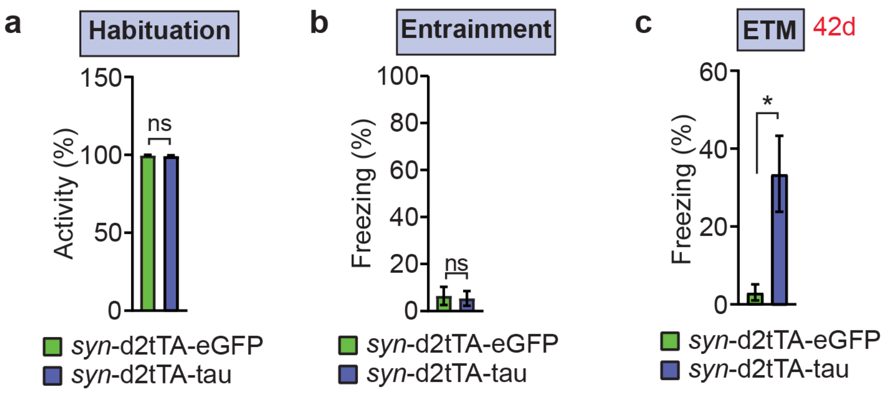

**Additional parameters of fear conditioning in mice with controlled neuronal tau expression**

(**a**) Locomotion during habituation, (**b**) freezing during entrainment, was comparable between *tau*$^{-/-}$ mice injected with AAV *syn1*-d2tTA-tau (n = 15) and *tau*$^{-/-}$ mice injected AAV *syn1*-d2tTA-eGFP (n = 16) with induced hippocampal eGFP or tau expression during entrainment (*see Fig. 2j*), which supports that short-term tau expression did not result in changes in trait or state anxiety.

(**c**) Significantly better memory recall at 42 days post-entrainment in *tau*$^{-/-}$ mice injected with AAV *syn1*-d2tTA-tau (n = 10) and *tau*$^{-/-}$ mice injected with AAV *syn1*-d2tTA-eGFP (n = 9) with induced hippocampal eGFP or tau expression during entrainment.

Statistical comparisons are performed using unpaired Student's t-test; *$p<0.05$; ns, not significant. Data are presented as mean ± S.E.M.

**Supplementary Figure 7**

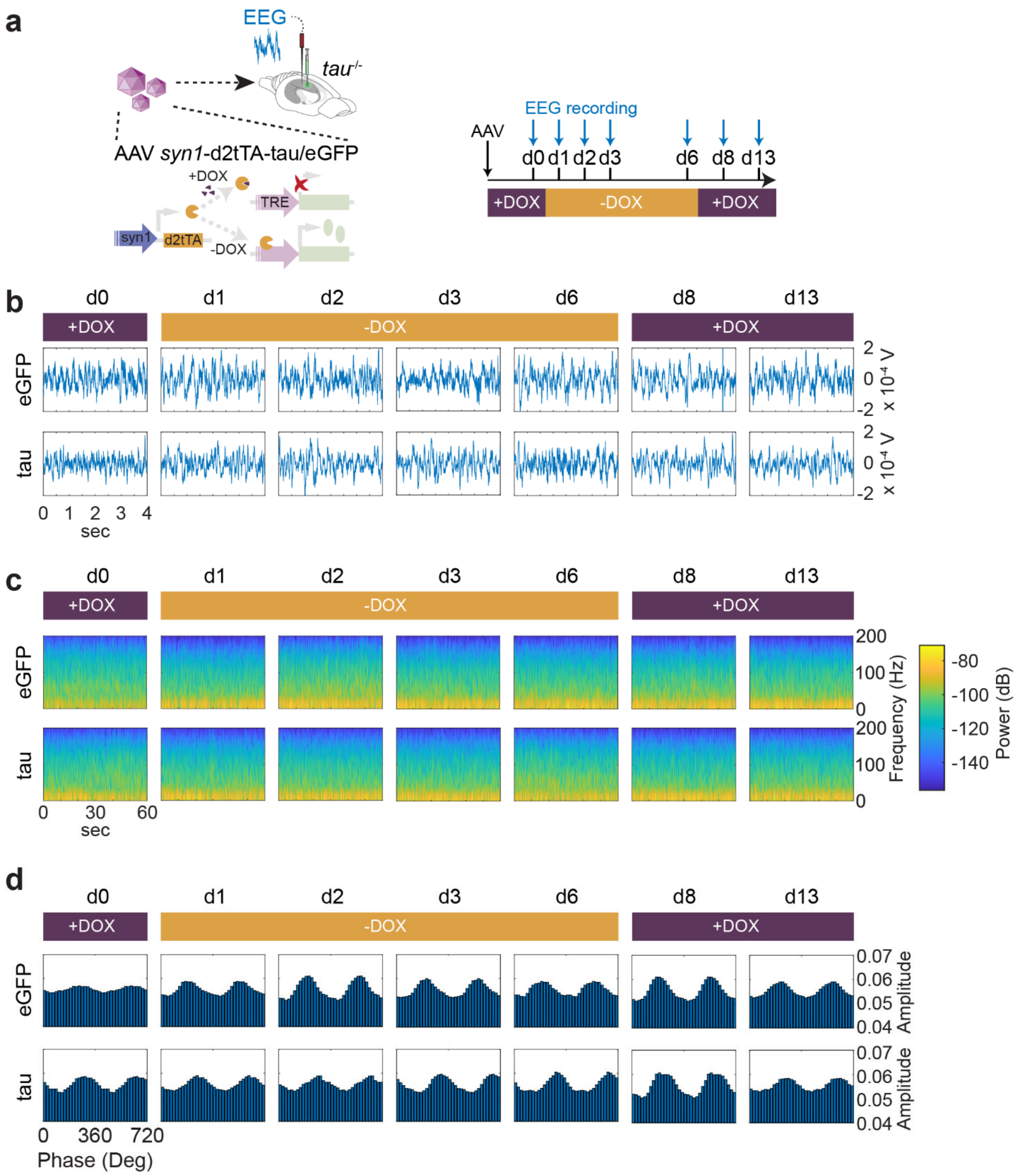

**Controlled transient neuron-specific tau expression does not alter activity, spectral distribution, and cross-frequency coupling in hippocampal networks.**

(**a**) Schematic of hippocampal EEG recording and doxycycline (DOX) diet feeding schedule for regulation of expression from AAV *syn1*-d2tTA-eGFP and -tau vectors injected into the hippocampus of *tau*$^{-/-}$ mice. Expression was induced by removal of DOX diet and placement of mice on regular chow.

Viral transgene expression was suppressed by refeeding with DOX diet. Unilateral hippocampal recording electrodes connected to wireless transmitters were implanted on the day of AAV injection. Mice were left on DOX diet for complete recovery from surgical procedure before recording.

(**b**) Representative hippocampal EEG traces (4 seconds) at indicated recording days from *tau*$^{-/-}$ mice injected with indicated *syn1*-d2tTA AAVs expressing either eGFP or tau.

(**c**) Spectrogram (dB) analysis for recordings represented in b. Epochs for recordings equivalent of 3 x 60 second bins (at 1,000 Hz sampling rate) were averaged.

(**d**) Cross-frequency coupling (Phase-Amplitude modulation plots) of 90s bins for recordings during expression schedule in a. Mean amplitude distribution over phase bins (20°) is shown over two theta phase cycles.

**Supplementary Figure 8**

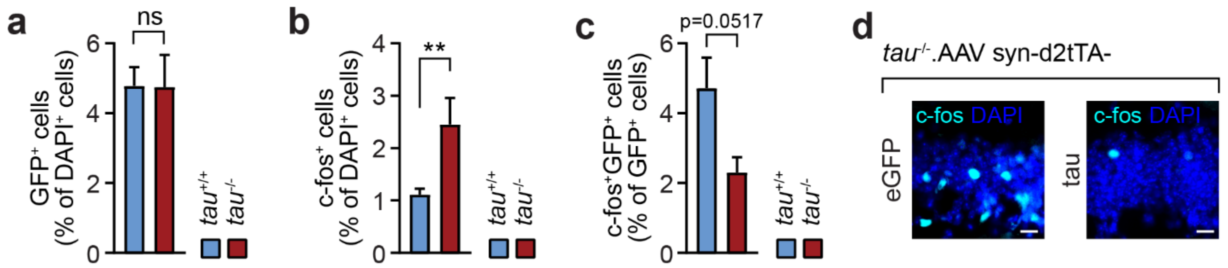

**Tau governs active neuron recruitment to the engram**

(**a**) Quantification of dentate gyrus (DG) cells associated with the engram (labelled with RAM-driven eGFP expression) in $tau^{+/+}$ and $tau^{-/-}$ mice during fear memory encoding. (n=6-7) Levels of $eGFP^{+}/DAPI^{+}$ cells indicated comparable engram labelling in both genotypes.

(**b**) Quantification of c-$fos^{+}$ DG cells in $tau^{+/+}$ and $tau^{-/-}$ mice during fear memory encoding. (n=6-7)

(**c**) Quantification of $eGFP^{+}$c-$fos^{+}$ double-positive cells relative to $eGFP^{+}$ engram cells in $tau^{+/+}$ and $tau^{-/-}$ mice during fear memory encoding. (n=6-7)

(**d**) Micrographs of c-$fos^{+}$ cells in hippocampal DG subfield of $tau^{-/-}$ mice expressing eGFP or tau during fear memory encoding ($tau^{-/-}$.AAV syn-d2tTA-eGFP/tau). DAPI, nuclei; scale bar, 20 µm (n=6-9)

Statistical comparisons are performed using Student's t-test; **$p<0.01$; ns, not significant. Data are presented as mean ± S.E.M.

**Supplementary Figure 9**

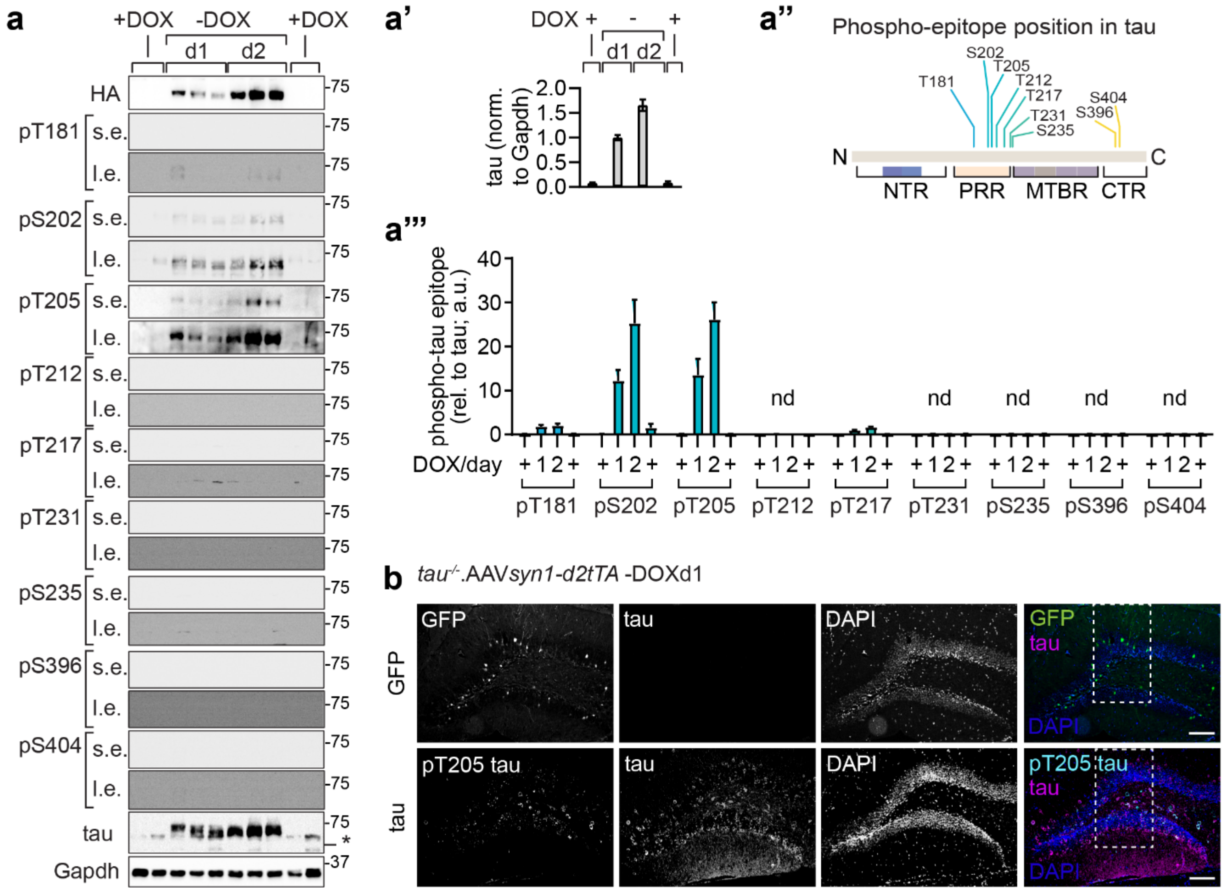

**Tau expressed during spatial acquisition only is phosphorylated at specific epitopes**

(**a**) Immunoblots of hippocampal lysates from tau-/- mice injected with AAV *syn1*-d2tTA-tau at indicated time points (d1-2) after removal of DOX-containing diet (-DOX) and start of spatial acquisition and at prior and subsequent suppression of tau by DOX-containing diet (+DOX). Immunoblots were probed for indicated phospho-epitopes of tau, total tau and Gapdh as loading control. Short (s.e.) and long exposure (l.e.) are shown for each antibody detection. *, unspecific band.

(**a'**) Quantification of hippocampal tau levels normalized to Gapdh loading control and relative to d1 of -DOX in *tau*$^{-/-}$ mice injected with AAV *syn1*-d2tTA-tau at indicated time points (d1-2) after removal of DOX-containing diet (-DOX) and start of spatial acquisition.

(**a''**) Position of phospho-epitopes detected in a relative to the primary sequence of tau. NTR, N-terminal region, including 2 facultative exons; PRR, proline-rich region; MTBR, microtubule-binding repeats; CTR, C-terminal region.

(**a'''**) Quantification of immunoblot signals for specific tau phospho-epitopes relative to and normalised to total tau signal. nd, not detectable

(**b**) Immunofluorescence labelling of hippocampal sections from tau-/- mice injected with AAV *syn1*-d2tTA-tau or -eGFP at day 1 (d1) after removal of DOX-containing diet (-DOX) and start of spatial acquisition. Scale bar, 200 µm.

Statistical comparisons are performed using ANOVA; $^{***}p<0.001$, $^{**}p<0.01$, $^{*}p<0.05$; ns, not significant. Data are presented as mean ± S.E.M.

**Supplementary Figure 10**

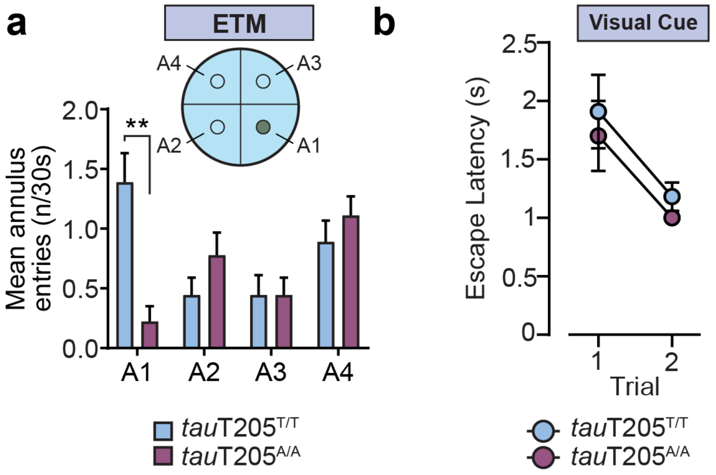

**Additional MWM analysis in *tau*T205T/T and *tau*T205A/A mice for spatial search strategy and extended-term memory**

(**a**) Mean annuli entries during extended-term memory (ETM) probe trial 4 months post-spatial acquisition towards annulus A1 (n=18-20).

(**b**) Visual cued trials confirm acuity in both *tau*T205T/T and *tau*T205A/A mice.

Statistical comparisons are performed using ANOVA; $**p<0.01$; ns, not significant. Data are presented as mean ± S.E.M.

**Supplementary Figure 11**

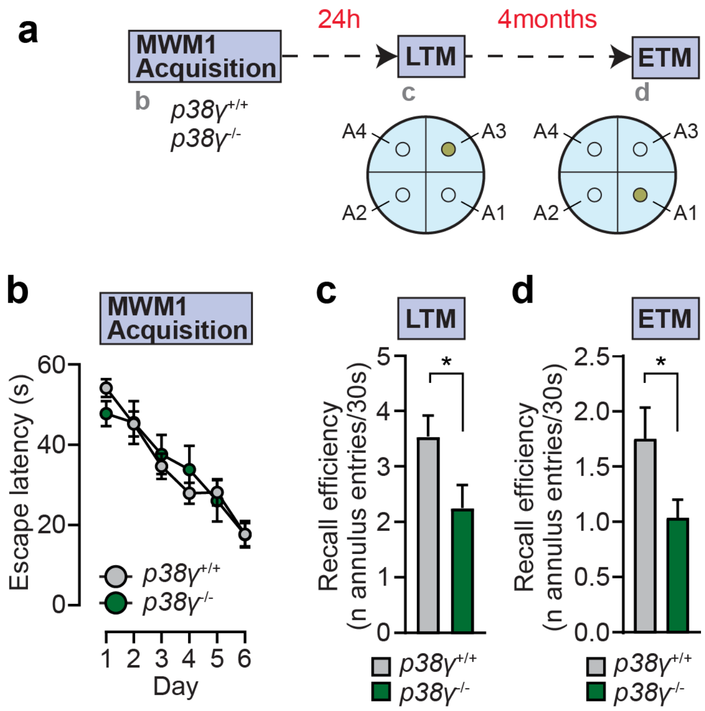

**MWM analysis in mice with gene deletion of MAP kinase *p38γ* (*Mapk12*)**

(**a**) Schematic of MWM to assess long-term (LTM) and remote (ETM) spatial memory in *p38γ* wildtype (*p38γ*$^{+/+}$) and *p38γ* knockout (*p38γ*$^{-/-}$) mice at 24 hour and 4 months post-spatial acquisition, respectively. Note that spatial acquisition for 24 hour recall (LTM) was to annulus A3, whereas acquisition for 4 months recall (ETM) was towards annulus A1 as target.

(**b**) Spatial acquisition curve (n=10-12) in *p38γ*$^{+/+}$ and *p38γ*$^{-/-}$ mice.

(**c**) Long-term memory (LTM) recall in *p38γ*$^{+/+}$ and *p38γ*$^{-/-}$ mice at 24 hour post-spatial acquisition. Recall efficiency is represented as mean annulus A3 entries per 30s of probe trial. Note that significant difference in spatial LTM is potentially due to impact of p38γ deletion on cognitive flexibility rather than annulus-specific effects.

(**d**) Extended-term memory (ETM) recall in *p38γ*$^{+/+}$ and *p38γ*$^{-/-}$ mice at 4 months post-spatial acquisition. Recall efficiency is represented as mean annulus A1 entries per 30s of probe trial.

Statistical comparisons are performed using ANOVA (b) or Student's t-test (c, d); *$p<0.05$; ns, not significant. Data are presented as mean ± S.E.M.

**Supplementary Figure 12**

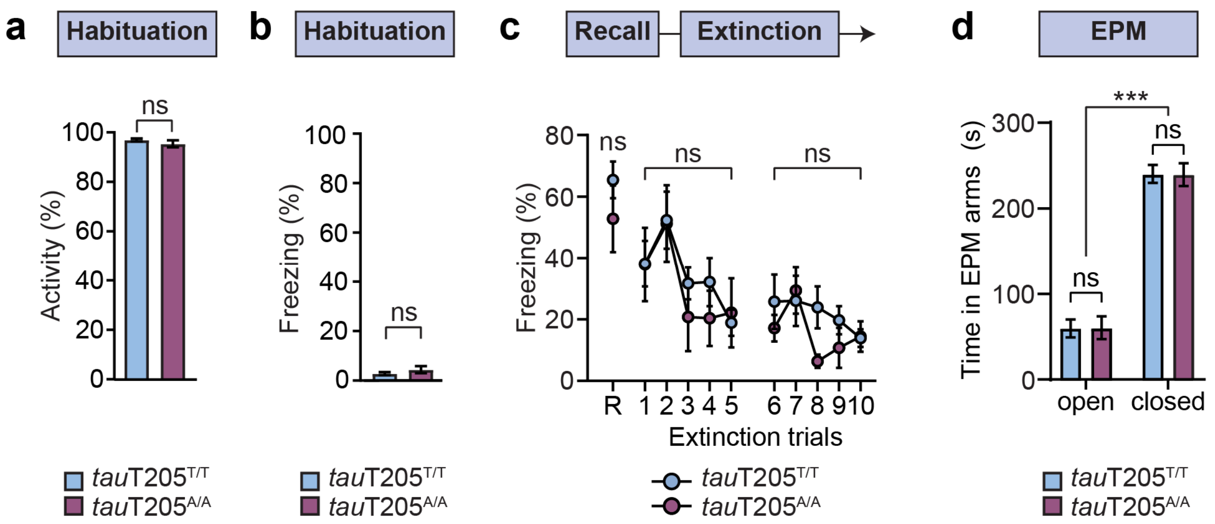

**Additional CFC analysis in mice with gene edited Threonine-205 phospho-site in tau**

(**a**) Activity recorded during habituation phase of CFC schedule in *tau*T205T/T and *tau*T205A/A mice. No difference was observed indicating comparable trait anxiety and general activity levels in both genotypes. (n=17-23) ns, not significant (Student's t-test).

(**b**) Spontaneous freezing during habituation phase in CFC schedule in *tau*T205T/T and *tau*T205A/A mice. No difference was observed indicating comparable trait anxiety. (n=17-23) ns, not significant (Student's t-test).

(**c**) Extinction of fear memory in *tau*T205T/T and *tau*T205A/A mice entrained by cued fear conditioning (n=8-12). Mice were exposed to an unpaired CS (2 min) in 5 consecutive trials on 2 consecutive days. Freezing response was recorded for initial recall of fear memory and each extinction trial. Note that *tau*T205A/A mice show comparable freezing response during extinction trials as *tau*T205T/T mice, precluding a role of tau T205 phosphorylation in active memory extinction.

(**d**) Spontaneous occupancy of open and closed arms of the elevated plus maze (EPM) for *tau*T205T/T and *tau*T205A/A mice. (n=8-12)

Statistical comparisons are performed using ANOVA (c, d) or Student's t-test (a, b); ***$p<0.001$; ns, not significant. Data are presented as mean ± S.E.M.

**Supplementary Figure 13**

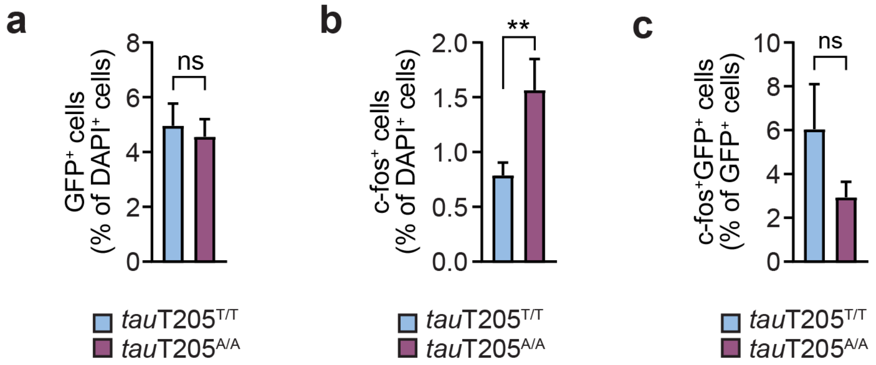

**T205 tau phosphorylation alters abundance and recruitment of active neurons to the engram**

(**a**) Quantification of dentate gyrus (DG) cells associated with the engram (labelled with RAM-driven eGFP expression) in *tau*T205T/T and *tau*T205A/A mice during fear memory encoding. (n=5) Levels of $eGFP^+/DAPI^+$ cells indicated comparable engram labelling in both genotypes.

(**b**) Quantification of c-fos$^+$ DG cells in *tau*T205T/T and *tau*T205A/A mice during fear memory encoding. (n=5)

(**c**) Quantification of $eGFP^+$c-fos$^+$ cells relative to $eGFP^+$ engram cells in *tau*T205T/T and *tau*T205A/A mice during fear memory encoding. (n=5)

Statistical comparisons are performed using Student's t-test; **$p<0.01$; ns, not significant. Data are presented as mean ± S.E.M.

**Supplementary Figure 14**

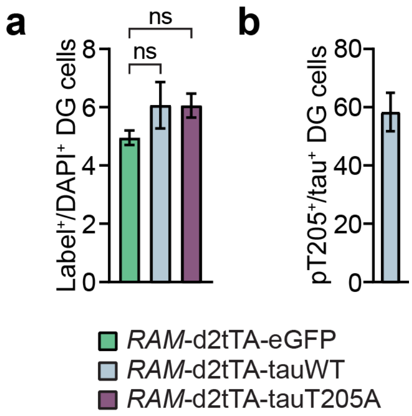

**Tau expressed in engram ensembles during memory acquisition is phosphorylated**

(**a**) Percentage of hippocampal dentate gyrus (DG) cells labelled with eGFP, tauWT or tauT205A by activity-dependent AAVs (RAM-d2tTA) on day 1 after DOX removal and spatial acquisition in the Morris water maze.

(**b**) Percentage of DG cells activity-dependently labelled with tauWT and positive for pT205 tau.

Statistical comparisons are performed using ANOVA; ns, not significant. Data are presented as mean ± S.E.M.

**Supplementary Figure 15**

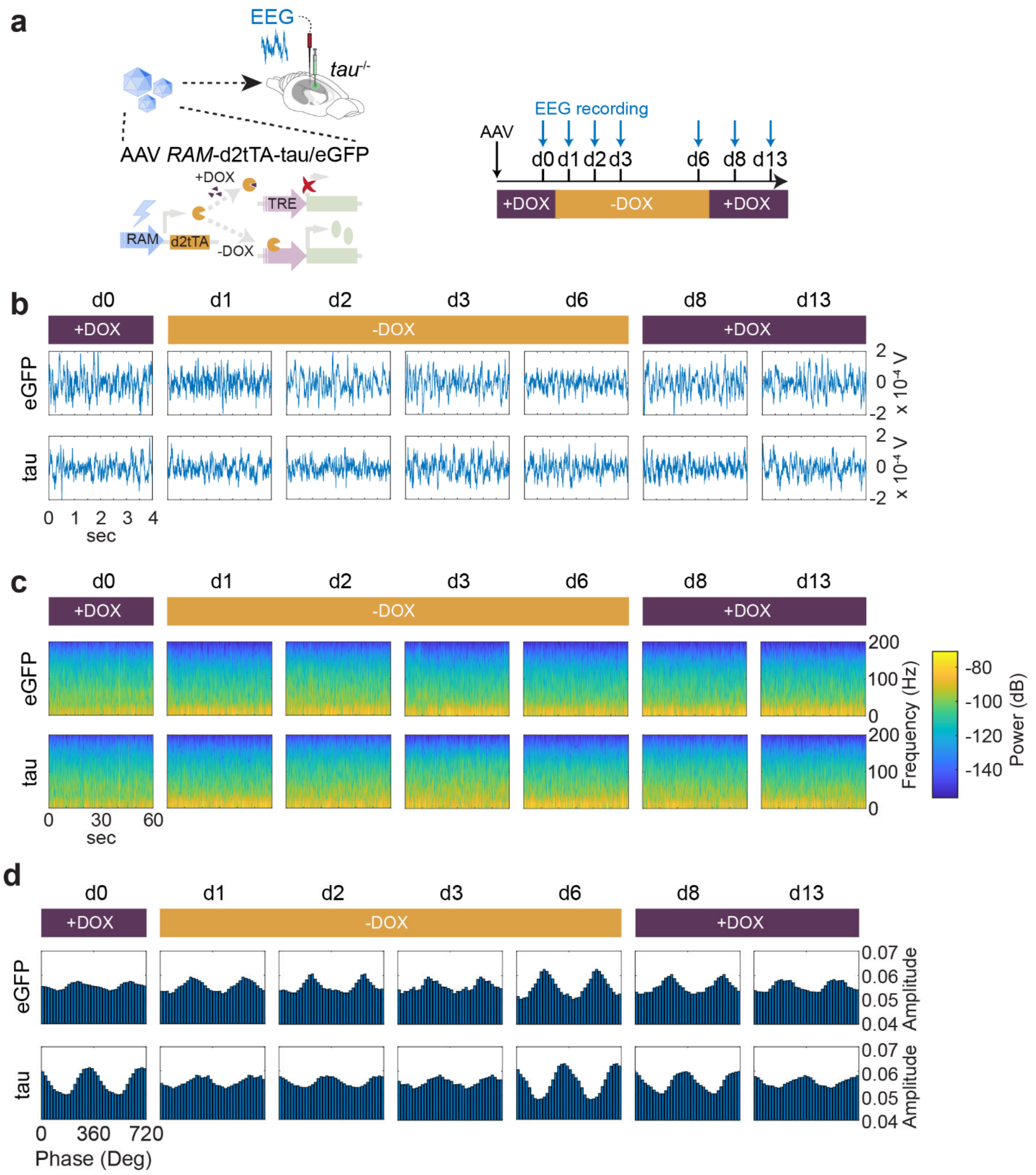

**Controlled transient expression of tau in active neurons does not alter activity, spectral distribution, and cross-frequency coupling in hippocampal local networks**

(**a**) Schematic of hippocampal EEG recording and doxycycline (DOX) diet feeding schedule for regulation of expression from activity-dependent AAV *RAM*-d2tTA-eGFP and -tau vectors injected into the hippocampus of *tau*$^{-/-}$ mice. Expression was induced by removal of DOX diet and placement of mice on regular chow. Viral transgene expression was suppressed by refeeding with DOX diet. Unilateral hippocampal recording electrodes connected to wireless transmitters were implanted on the day of AAV injection. Mice were left on DOX diet for complete recovery from surgical procedure before recording.

(**b**) Representative hippocampal EEG traces (4 seconds) at indicated recording days from $tau^{-/-}$ mice injected with indicated *RAM*-d2tTA AAVs expressing either eGFP or tau.

(**c**) Spectrogram (dB) analysis for recordings represented in b. Epochs for recordings equivalent of 3 x 60 second bins (at 1,000 Hz sampling rate) were averaged.

(**d**) Cross-frequency coupling (Phase-Amplitude modulation plots) of 90s bins for recordings during expression schedule in a. Mean amplitude distribution over phase bins (20°) is shown over two theta phase cycles.

(c) Cross-frequency coupling analysis with modulation index at different stages of suppression and induction of activity-dependent tau expression.

**Supplementary Figure 16**

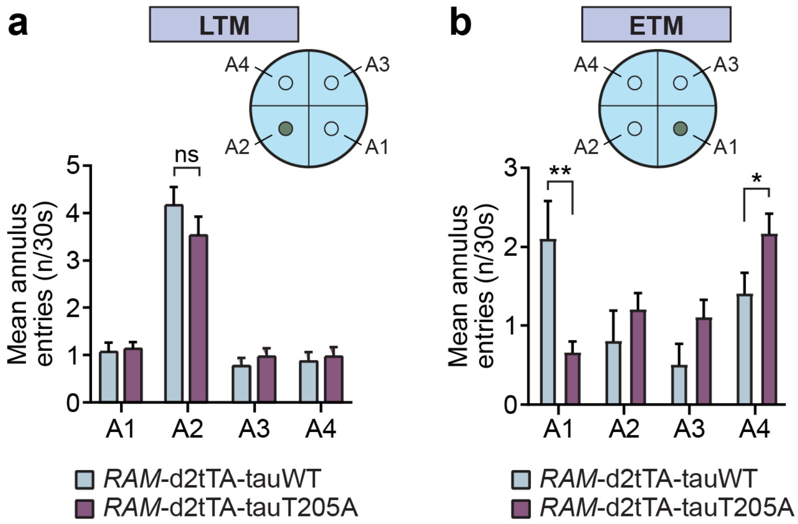

**Additional MWM analysis in mice with engram-restricted tauWT or tauT205A expression during memory encoding**

(**a**) Long-term memory (LTM) recall for spatial acquisition towards annulus A2 in *tau*$^{-/-}$ mice hippocampally injected with indicated activity-dependent *RAM*-d2tTA AAVs expressing either tauWT or tauT205A at 24 hour post-spatial acquisition (n=14-15). Recall efficiency is represented as mean annulus entries per 30s of probe trial.

(**b**) Extended-term memory (ETM) recall for spatial acquisition towards annulus A1 in *tau*$^{-/-}$ mice hippocampally injected with indicated activity-dependent *RAM*-d2tTA AAVs expressing either tauWT or tauT205A at 4 months post-spatial acquisition (n=14-15). Recall efficiency is represented as mean annulus entries per 30s of probe trial.

Statistical comparisons are performed using ANOVA; $^{**}p<0.01$, $^{*}p<0.05$; ns, not significant. Data are presented as mean ± S.E.M.

**Supplementary Figure 17**

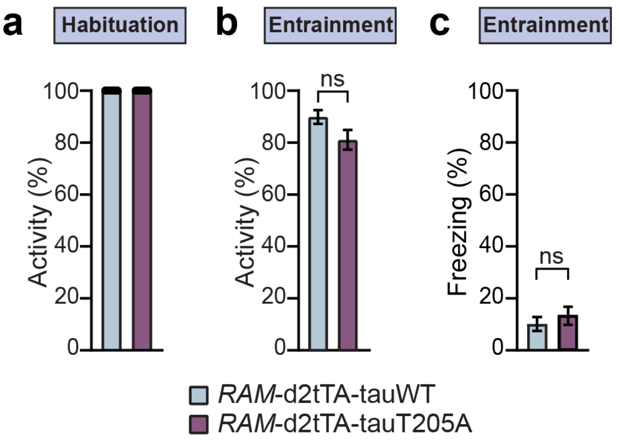

**Comparable spontaneous activity and freezing in tau-/- mice upon engram cell-specific expression of tauWT and tauT205A**

(**a**) Activity recorded during habituation phase of cued fear conditioning schedule in *tau*$^{-/-}$ mice hippocampally injected with indicated activity-dependent *RAM*-d2tTA AAVs expressing either tauWT or tauT205A. No difference was observed indicating comparable trait anxiety and general activity levels in both genotypes. (n=16-18)

(**b**) Activity recorded during entrainment phase of cued fear conditioning schedule in *tau*$^{-/-}$ mice hippocampally injected with indicated activity-dependent *RAM*-d2tTA AAVs expressing either tauWT or tauT205A. No difference was observed indicating comparable trait anxiety and general activity levels in both genotypes. (n=16-18)

(**c**) Spontaneous freezing during entrainment phase in CFC schedule in *tau*$^{-/-}$ mice injected with *RAM*-d2tTA AAVs expressing either tauWT or tauT205A. No difference was observed indicating comparable trait anxiety. (n=16-18)

Statistical comparisons are performed using Student's t-test; ns, not significant. Data are presented as mean ± S.E.M.

**Supplementary Figure 18**

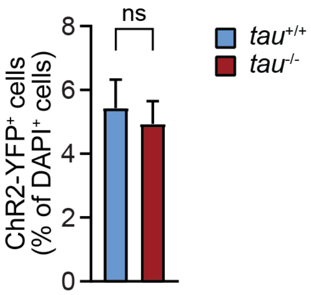

**Labelling for optogenetic engram activation in *tau*-deficient mice on DOX.**

(**a**) DG sections (*n* = 6-7 mice per group) revealed comparable ChR2–eYFP labelling induced at CFC entrainment, consistent with the previously established engram tagging strategy (Ref. 5, 70).

Statistical comparison was performed using Student's t-test; ns, not significant. Data are presented as mean ± S.E.M.
